# The gene-like promoter and transcription of LTR retrotransposons

**DOI:** 10.64898/2026.05.08.723830

**Authors:** Caleb Gooden, Xingli Li, Isabella Walter, Shujun Ou

## Abstract

Transcriptional regulation is one of the fundamental approaches for young plants to cope with environmental fluctuations and maintain active development. The transposable element (TE) subclass long terminal-repeat retrotransposons (LTR-RTs) can act as additional regulators for genes through enhancer and promoter activity, but their promoters, transcription initiation, and contributions during maize development remain uncharacterized. Here, we developed IsoClassifier to resolve the transcription start site (TSS) and RNA isoforms of LTR-RTs based on long-read transcriptomics, delineating LTR U3 regions as the native promoter and enhancer of LTR-RTs. We reveal conserved motifs associated with core promoter activity in transcribed LTR-RTs that are highly comparable to gene promoters. Further, we found that LTR-RT transcription in maize was dominated by spliced, long non-coding RNA. Finally, a genome-wide coexpression analysis revealed that LTR-RTs are transcribed as hub-like elements in coexpression networks, suggesting important roles in gene regulation. We conclude that LTR-RTs have similar promoter compositions to gene promoters and likely share similar transcription regulation programs.

## Main

Retrotransposons are mobile genomic segments found in most eukaryotic organisms. In plants, long terminal-repeat retrotransposons (LTR-RTs) dominate the genomic content and are transposed using RNA intermediates^1^. LTR-RTs are characterized by an internal coding region flanked by identical, noncoding LTRs. Recent work demonstrates that LTR-RTs may provide *cis*-regulatory elements for gene regulation, which could drive phenotypic variation and tissue-specific expression^2–10^. LTR-RTs comprise 75% of the maize genome and represent an underexplored regulatory reservoir. Such regulatory information is usually found in the LTR region of the element that harbors the putative promoter region of LTR-RTs. However, the precise location or characteristics of LTR promoters are mostly unknown due to the divergence of LTR-RT families^1,11–13^.

The LTR region can be further divided into subregions termed U3, R, and U5. The U3 subregion contains putative promoter and enhancer features, which could act as *cis*-regulators for nearby genes^14^. Examples include LTR-RT insertions enhancing *MdMYB1* expression in apples^4^ and *Ruby* expression in blood oranges^8^, both resulting in darker fruit coloration. However, these studies focused on the entire LTR-RT element and their effects on nearby gene transcription. Identifying and characterizing promoter regions of LTR-RTs will address a knowledge gap in LTR-driven *cis*-regulatory activities, linking proximity effects of mobile element insertions to molecular mechanisms.

A well-studied case of the U3 promoter is the Human Immuno-deficiency Virus (HIV). HIV is a LTR-RT-like retrovirus that can similarly recruit RNA polymerase II (RNAP-II) of the host for its own transcription and replication. HIV’s U3 region has been noted to harbor binding sites for modulatory transcription factors and contain core promoter motifs like the TATA box^15^. The motif sites are used temporally by the virus to control transcription levels depending on host and cell state. For example, an activator protein 1 (AP-1) binding site in the U3 regions enables establishment of HIV latent infection programs^15,16^, while a specificity protein 1 (Sp1) binding site can work in tandem with the TATA box to direct basal transcription or with inducible transcription factors to organize active infection^15,17^. Currently, there is a lack of comparable knowledge for LTR retrotransposons in plants. Analysis of LTR-RT composition has been attempted but ultimately rendered incomplete due to shortfalls of short-read sequencing methods in resolving repetitive LTR-RT loci^3,13,18^. The boundaries and features of the U3 region responsible for recruiting transcription components and contributing to cellular gene regulation in plants thus remains largely uncharacterized.

We hypothesized that LTR U3 promoters harbor conserved features responsible for their ubiquitous ability to be transcribed by host RNAP-II despite profound sequence divergence^1,11–13^. Advancements of long-read sequencing, like Oxford Nanopore Technology’s cDNA sequencing, have greatly improved the accuracy in distinguishing the expressions of highly repetitive LTR-RTs and their isoforms^19–22^. The work presented here takes advantage of long-read sequencing technologies to identify LTR-RT promoters and explore their genetic and epigenetic features contributing to transcription initiation. We compared the similarity of LTR-RT promoters to genes and identified links to regulatory networks of maize development.

## Results

### Long-read sequencing captures full-length LTR-RT transcripts

Since the promoter of LTR retrotransposons is located in the U3 subregion of the terminal repeat, we first identified transcription start sites (TSS) to mark the boundary of U3 with the R subregion. We developed two bioinformatics pipelines: 1) “AccuMap” for long-read transcriptome stranding and alignment, and 2) “IsoClassifier” to identify transcribed LTR-RT loci and predict their TSS based on alignment consensus (**Fig. 1A,D**).

**Figure 1.**
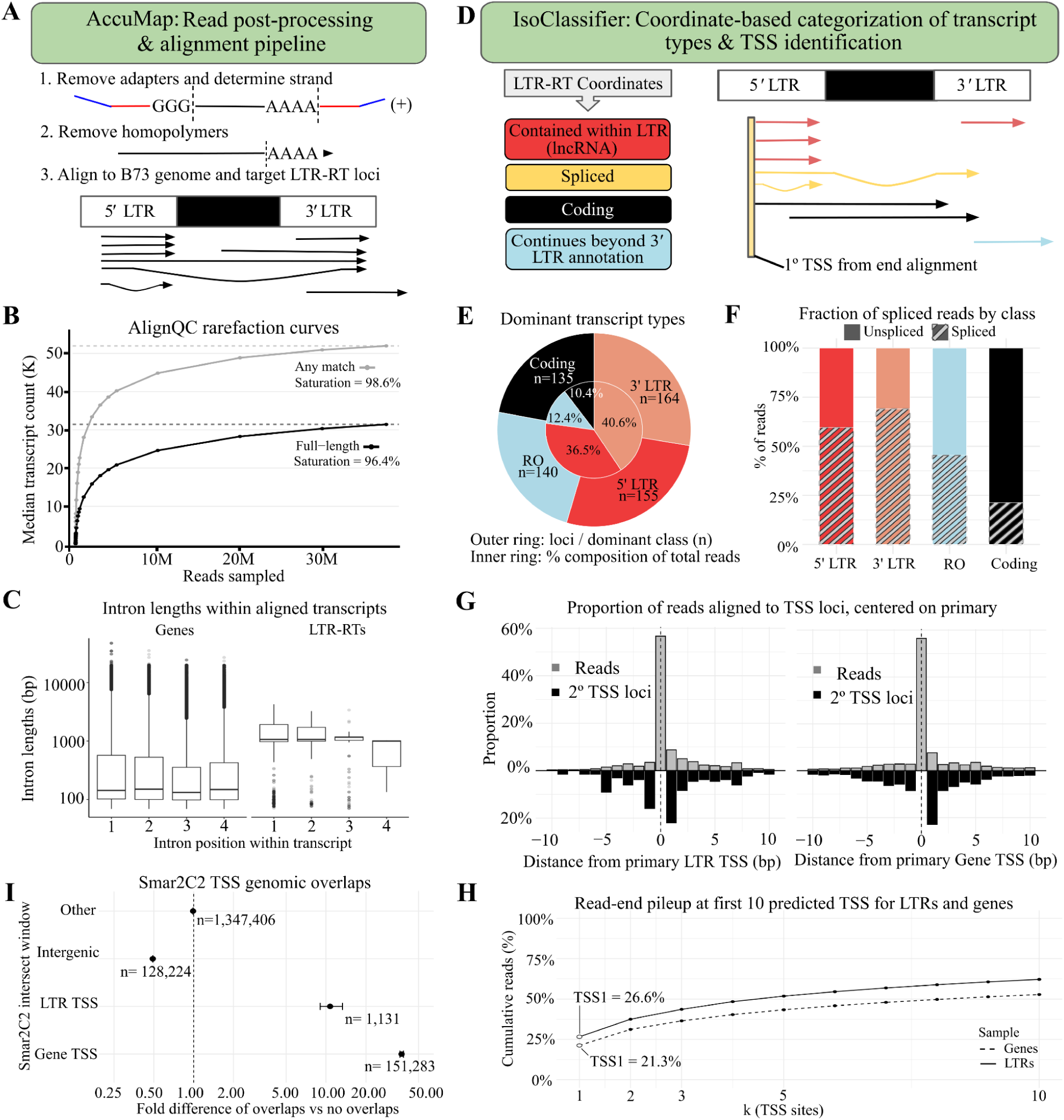
Long-read RNA-seq accurately identifies LTR-RT transcription start sites (TSS) and distinguishes transcription isoforms. **A)** The AccuMap pipeline uses sequencing adaptors and poly(A) tail locations for transcript orienting. **B)** AlignQC rarefaction subsampling on Nanopore long-read transcriptome data used in this study for any-match (grey) and full-length (black) alignments. Dashed lines mark the observed maxima. The Michaelis-Menten model was used to estimate the degree of saturation. **C)** Log-transformed length distributions of the first four introns in transcribed genes and LTR-RTs. **D)** IsoClassifier determined LTR-RT transcript isoforms based on their relative locations to the LTR-RT. E) LTR-RT isoform abundances on the locus level (outer ring) and transcript level (inner ring). RO = LTR-RT transcription continues beyond 3′ LTR (read out); Coding = Transcripts spanning the LTR-RT internal region; 5′ LTR and 3′ LTR = Transcripts contained in the 5′ and 3′ LTR, respectively; **F)** Fraction of spliced transcripts among different LTR-RT transcript classes. G) Read pileup (grey) of 5′ read-ends ± 10bp of the primary (1°) TSS relative to all reads aligned in this region; the distribution of secondary (2°) TSS predictions (black) relative to the primary TSS. **H)** Cumulative 5′ read end contributions to TSS predictions of LTR-RTs and genes. The k-value represents incremental alternative TSS following the primary prediction (k = 1). I) Permutation enrichment test for IsoClassifier TSS primary prediction within ±100bp of Smar2C2 sites. The genomic background (Other) is used as the baseline. Horizontal bars represent the 95% confidence interval.

AccuMap takes raw sequencing results of Nanopore long-read cDNA as input and uses PyChopper^23^ to remove sequencing adapters and orient read strands, cutadapt^24^ to remove poly(A) and poly(T) tails, and minimap2 (ref ^25^) for splice-aware alignments. We generated three cDNA libraries from the upperground tissues of maize B73 seedlings 21 days after sowing (DAS), which were sequenced on a Nanopore PromethION machine. This yielded a total of 49.6 million raw reads. Using AccuMap, we eliminated 10.1 million reads that had too little information to be demultiplexed or strand-oriented, retaining 39.5 million reads which were aligned to the B73v5 genome^26^. Of these, 37.1 million (94.1%) reads were mapped as primary alignments. We incorporated low-depth PacBio IsoSeq data^21^ from embryos, endosperm, pollen, tassels, roots, and ears with our Nanopore reads for additional identification of LTR-RT isoforms. This yielded a final library of 37.9 million primary alignments (**Table S1**).

Rarefaction assessment of isoform discovery using AlignQC^27^ showed that the full-length transcript count of the Nanopore transcriptome was saturated with 38 million reads. This suggested that our data had sufficient depth to cover rare and lowly expressed isoforms (**Fig. 1B; Table S1**), providing confidence in revealing a comprehensive view of LTR-RT transcription in the maize B73 genome. We further applied a minimum MapQ score of 30 to filter out low-quality alignments, resulting in a total of 34 million primary alignments (**Table S1, Fig. S1**) where 94.2% of these reads overlapped LTR-RTs and/or genes. Globally, 1.3% of primary aligned transcripts (0.4 million) overlapped with LTR-RTs, and 92.3% of LTR-RT transcripts overlapped the Ty3 superfamily or LTR-RTs that lack any coding domain for classification (**Table S1**). To study LTR-initiated transcriptions, we only retained transcripts that were initiated from the LTR region of structurally intact LTR-RTs, producing a higher-quality set of 13,012 transcripts: 1,139 from the Ty1 superfamily, 3,191 from the Ty3 superfamily, and 8,682 from LTR-RTs of unknown classification. Despite sufficient total read coverage (**Fig. 1B)**, the read-count distribution of LTR-RTs showed an excess of low-coverage loci suggesting transcriptional noise. We fit a segmented regression to each sample (**Table S2**) and used a sigmoid probability function to confidently identify 594 LTR-RTs yielding RNA transcripts for further study.

We observed abundant transcripts contained within the LTR region which carried a previously reported splicing pattern with unclear biological significance^22^. These spliced LTR transcripts often displayed three exons: a short middle exon flanked by two longer and separated by long introns (**Fig. 1C; Fig. S2**). The average LTR exon lengths (**Fig. S3A**) and intron lengths (**Fig. 1C**) were 245 and 1303 bp, respectively, longer than gene exons (217 bp) and introns (469 bp). We also validated canonical splice junctions for all gene and LTR transcripts reported as spliced and observed over 98% read support (**Fig. S3B**). Examination of read 3′ clipping and cleavage sites (poly(A) trimming by cutadapt) showed that LTR-initiated transcripts were fully aligned to the genome (**Fig. S3C-F)**, supporting them as valid transcription events with confident origin tracing.

Previous work demonstrated that LTR-RTs can produce distinct transcript isoforms with different implications for function^28^. Based on how LTR-initiated transcripts aligned to the genome, we developed a module in IsoClassifier to separate LTR-RT transcripts into four categories (**Fig. 1D,E**): “spliced” (reads that have evidence of splicing and splice junctions), “LTR-contained” (read starts and ends within the LTR region), “LTR-coding” (read covers the 5’ LTR and coding region and ends in the 3′ LTR), and “read-out” (begins in an LTR region and extends beyond the 3′ end of the element). The “spliced” classification is a status that is applied to the other three categories. Transcripts restricted in LTR regions were found in 360 out of 594 LTR-RTs and constituted the dominant isoform of 319 loci (53.7% of all loci; **Fig. 1E**). These “LTR-containted” transcripts had a minimal and average length of 81 and 666 bp, respectively, which follows the general definition of long-noncoding RNA (lncRNA)^29,30^. We found that 77.1% of all LTR-RT transcripts were non-coding LTR-contained isoforms, suggesting abundant lncRNA transcription (**Fig. 1E**). This result agrees with previous reports that between ∼43% and ∼57% of lncRNA transcripts originate from LTR-RTs^29,31^. The higher fraction of LTR-derived lncRNA detected in our study likely resulted from the improved resolution of highly repetitive sequences by long-read sequencing.

A per-locus overview found that ∼50% of transcribed 5′- and 3′-LTR loci produced at least one spliced transcript (**Fig. S4A**); 59.6% of 5′-LTR and 69.2% in 3′-LTR-contained reads exhibited splicing patterns (**Fig. 1F**) with a minimum combined exon length of 204 bp and average of 698 bp. We identified 178 LTR-RTs that predominantly produced spliced, noncoding transcripts (**Fig. S3, Fig. S4)**. We calculated the sequence identity between LTR regions using Kmer2LTR^32^ to approximate an element’s age, and observed that loci with spliced LTR-contained transcripts were significantly younger than those with non-spliced (Wilcoxon test, *p* < 9.4 × 10^-12^) and younger than those structurally-intact, silent LTR-RTs (Wilcoxon test, *p* = 1.0 × 10^-6^; **Fig. S4B,D,E**). We additionally discovered that LTR identity was weakly but positively correlated with the percentage of spliced reads (Spearman’s ρ = 0.4, *p* = 2.4 × 10^-15^) (**Fig. S4B**). A binomial regression confirmed that splicing frequency increased significantly with increasing LTR identity, and after correcting for overdispersion, the effect remained robust (*p* = 1.7 × 10^-4^; **Fig. S4C**).

### Accurate identification of LTR-RT TSS

The TSS of LTR-RTs serves as the boundary between the U3 and R subregions of the terminal repeat, so its identification can be used to delineate the LTR promoter (U3). Based on full-length cDNAs, we used the collective starting position of alignments to identify TSSs for LTR-RTs and define the U3/R boundary. This functionality was integrated into IsoClassifier and detects multiple TSS candidates based on transcript support. The top 10 TSS candidates of all transcribed LTR-RTs were cumulatively supported by over 62% of LTR-RT transcripts. Primary TSS candidates in LTR-RTs were supported by 26.6% of total transcripts (**Fig. 1H**). In comparison, the top 10 gene TSS candidates were supported by over 53% of gene transcripts, with the primary TSS candidates supported by 21.3% (**Fig. 1H**).

Alternative TSSs are biologically meaningful^3,21^ and our data suggest the prevalence of alternative TSS in both LTR-RTs and genes, with more than 50% of transcripts initiated from a 20-bp window around a predicted primary TSS (**Fig. 1G**). We observed more secondary TSS candidates appearing downstream of the predicted primary TSS (**Fig. 1G**), suggesting potential mRNA degradation or processing. Because reads from cDNA-based RNA sequencing are subject to end-degradation that can still lead to imprecise alignments after end-processing, the asymmetrical alternative TSS distributions are consistent with end biases inherent to RNA sequencing. However, read support for the primary TSS prediction is much stronger than that of alternative TSSs. To validate our pipeline, we compared predicted TSS to experimentally-determined TSSs from Smart-Seq2 Rolling Circle to Concatemeric Consensus (Smar2C2-seq)^33^ sequencing and Cap Analysis Gene Expression and deep sequencing (CAGE-seq)^34^ in maize B73 root and shoot tissues. Our TSS predictions in both genes and LTRs were highly colocalized with TSSs annotated by Smar2C2-seq and CAGE-seq (**Fig. S2**; **Fig. S5**). A permutation-based Monte-Carlo enrichment/depletion analysis revealed that Smar2C2 peaks are significantly concentrated near the IsoClassifier-predicted TSSs of both genes and LTR-RTs (*p* < 0.001), and depleted from intergenic regions (*p* < 0.001) when compared to background (**Fig. 1I**). Together, these data demonstrated that our method reliably predicted TSSs for both genes and LTR-RTs, allowing the identification of LTR U3 promoter boundaries.

### LTR-RT promoters share similar features with gene promoters

Determining the process of host RNAP-II recruitment for LTR-RT transcription was our next step in studying retrotransposons’ regulatory functions. We conducted a motif analysis to compare core promoter features of LTR-RTs with gene promoters. We developed “WindowScrubber,” a position-weight matrix (PWM) approach to locate pre-defined motifs within specified sequence ranges relative to a positional anchor, which is the primary TSS in our case. This method enabled us to locate motifs in each transcribed LTR-RT and compare motif distributions around the TSS for U3s that differed in size.

We used WindowScrubber to scan for canonical promoter motifs, including TATA boxes, pyrimidine patches (“Y-patches”), CCAAT boxes, downstream promoter elements (DPE, consensus sequence *5′-RGWCGTG-3′*), and initiator (Inr, consensus sequence *5′-PTCA+1NTPP-3′*) sequences^34,35,35–38^. Despite different numbers of transcribed loci, we found highly similar enrichment and positional distributions of core promoter motifs between LTR-RTs and genes (**Fig. 2B; Fig. S6**). Upstream TATA boxes appeared in 261 (43.9%) of LTRs and 7,995 (29.1%) of genes within 120 bp of the predicted primary TSS (**Fig. 2A,B**). Both genes and LTR-RTs displayed the same peak of TATA motifs at -32nts to their respective primary TSS. These results are within the canonical TATA box range for maize and agree with previous reports describing primary TATA peaks between -55nt and -23nt^39^.

**Figure 2.**
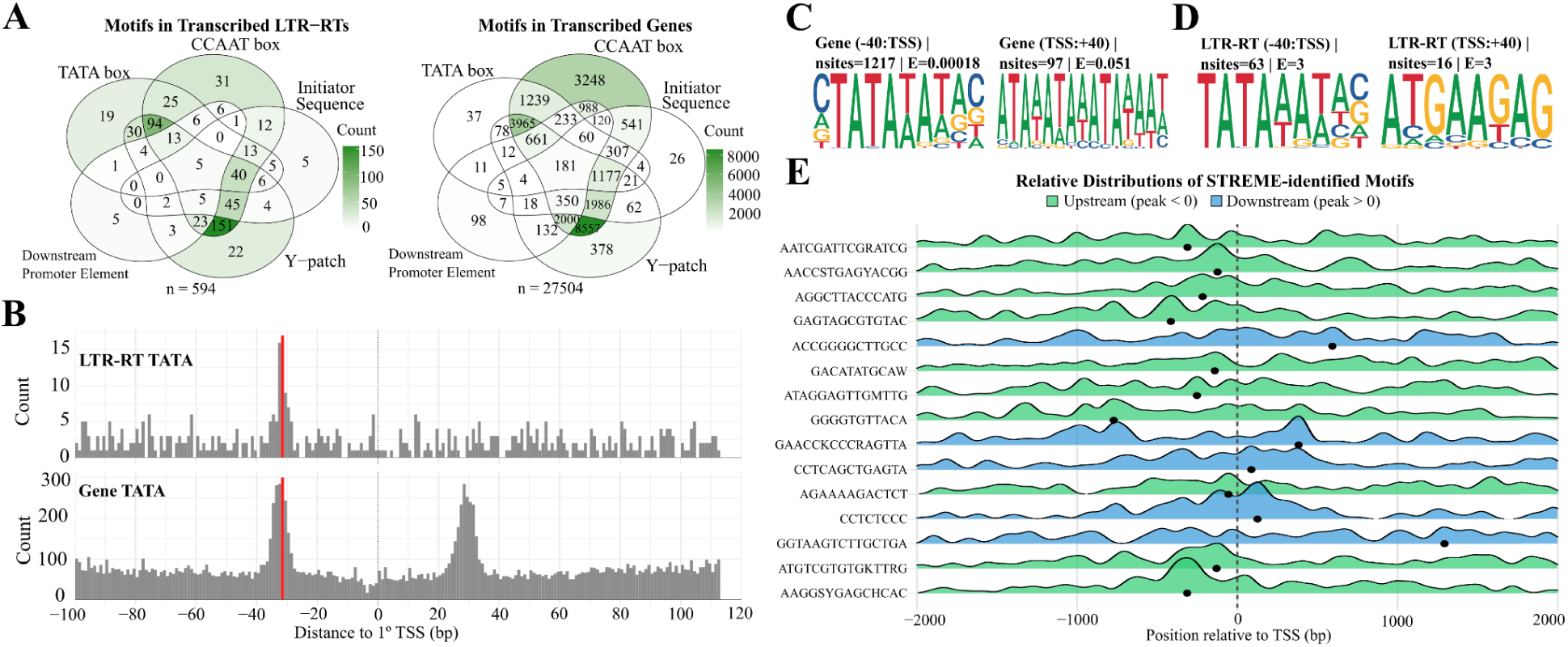
LTR-RTs share similar core promoter features with gene promoters. **A)** Composition of motifs found by WindowScrubber for gene and LTR-RT promoters of elements passing threshold filtering (594 LTR-RTs, 27,504 genes). **B)** TATA distribution relative to TSS for genes and LTR-RTs. Y axes show the number of loci with matches starting at position *x*. The primary TATA-box candidate is highlighted in red. **C)** Sequence logos of the two TATA peaks in genes. The canonical TATA box *5′-TATA(A/T)A(A/T)(A/G)-3′* is enriched upstream but not downstream of gene TSS. **D)** The canonical TATA box is discoverable upstream of LTR TSS but not downstream. **E)** Novel motifs (**Table S3**) are positionally enriched around the LTR TSS. The black dot at the base of each ridge represents that motif’s highest density in the region.

Genes displayed an additional secondary TATA peak at +28nts consisting of 8,837 (32.1%) loci, which had not previously been reported (**Fig. 2B**). Further motif scan by STREME^40^ revealed significant enrichment of the canonical TATA boxes upstream but not downstream of the predicted primary TSS (**Fig. 2C**). The canonical TATA box was also found upstream but not downstream of the LTR TSS, although instances were too low for significance (**Fig. 2D**). In summary, we report that the canonical TATA box was found in both gene and LTR-RT core promoters.

The 8-mer Y-patch motif^35,36^ appeared in 19,582 (71.2%) of genes and 447 (75.3%) of LTR-RTs within 100bp upstream of our primary TSS predictions. The Inr sequence is commonly found in promoters lacking a TATA box and overlapping the TSS^34,35^. Of the 4,869 genes and 143 LTR-RTs harboring an initiator overlapping the TSS in our dataset, 3,110 (63.9%) of the genes and 74 LTR-RTs (51.7%) lacked a primary TATA box. The DPE can work in conjunction with Inr downstream of the TSS and is found predominantly in promoters lacking a TATA box^35,37,38^. Scanning for DPEs uncovered 4,880 genes and 74 LTR-RTs carrying the motif, 745 and 13 respectively also contained an initiator sequence, and 3,713 and 45 respectively lacked a TATA box (**Fig. 2A**). The most abundant motif found in our features was the CCAAT-box transcription factor binding site^35,41^ in its canonical -140:-460bp range, matching 470 (79.1%) LTR-RTs and 25,613 (93.1%) genes. The highly similar core promoter motif compositions between LTR-RTs and genes provide strong evidence for their similar RNAP-II recruitment utility (**Fig. S6**).

Given their core promoter characteristics, we were also interested in transcription factor binding motifs (TFBMs) enriched in LTR promoters. We randomly selected a transcribed LTR-RT from each transcribed family (n = 256) to form a nonredundant set of promoter sequences. We searched *de novo* for TFBMs using STREME in these non-redundant LTR promoters and identified 15 candidate motifs (**Fig. 2E**). Using TOMTOM^42^ and the JASPAR non-redundant plant motifs database^43^, none confidently matched known TFBMs in plants, suggesting these elements may bind uncharacterized or highly divergent maize-specific transcription factors. With FIMO^44^, we found that most of the 15 motifs clustered near the primary LTR TSS (**Fig. 2E**). These data support a multi-factorial transcription regulatory system for LTR-RTs in maize similar to those seen in the LTRs of HIV^15–17^ and gene promoters^33–37,43^.

### LTR-RTs share similar spatial epigenetics and transcription profiles with genes

Since LTR-RTs are known to affect the transcriptional landscape, we further explored the spatial transcription of LTR-RTs from multiple tissues. The ONT and PacBio transcriptome data originated from pollen, embryos, endosperm, leaves, shoots, roots, ears, and tassels had highly uneven coverages. We also included high-depth RNA-seq data collected by Illumina NextSeq 550 in B73 tissues^26^, including 16 day-after-pollination (DAP) embryos, 16 DAP endosperm, 8 day-after-sowing (DAS) root, 8 DAS shoot, R1 anther, V11 leaves, V18 ears, and V18 tassels to supplement our current data pool (**Table S1**).

To better handle the complexity and variability of samples in our dataset, we developed the **De**tailed **C**haracterization of **L**ong-**T**erminal **R**epeats (DeCLTR) pipeline (**Fig. 3A**) as an alternative to Multi-Omics Factor Analysis (MOFA)^45^ and Neuroscience Multi-omic Archive (NeMO)^46^ for classification of expression. DeCLTR is a reduction method that makes approximations of global activity using probabilities of multi-omics data overlap as supporting evidence. Multi-omics data is compiled into a matrix with entries for each LTR-RT and gene and assigns an “Activity” category based on omics signals. In addition to adding transcriptomics and TSS validation like Smar2C2 and CAGE into the matrix, we examined CpG DNA methylation and histone modifications H3K4me3 and H3K27ac^47^. Trimethylation of H3K4 is most commonly associated with transcription initiation and elongation and usually overlaps a TSS^48,49^. Active enhancers and promoters are also commonly associated with H3K27 acetylation^49,50^. In contrast, CpG methylation is a modification linked to long-term transcriptional silencing, where hypomethylated CpG islands (stretches of CpGs) are commonly found in promoter regions^51^. These three -omics additions to our study further contextualized the LTR-RT transcriptional landscape.

**Figure 3.**
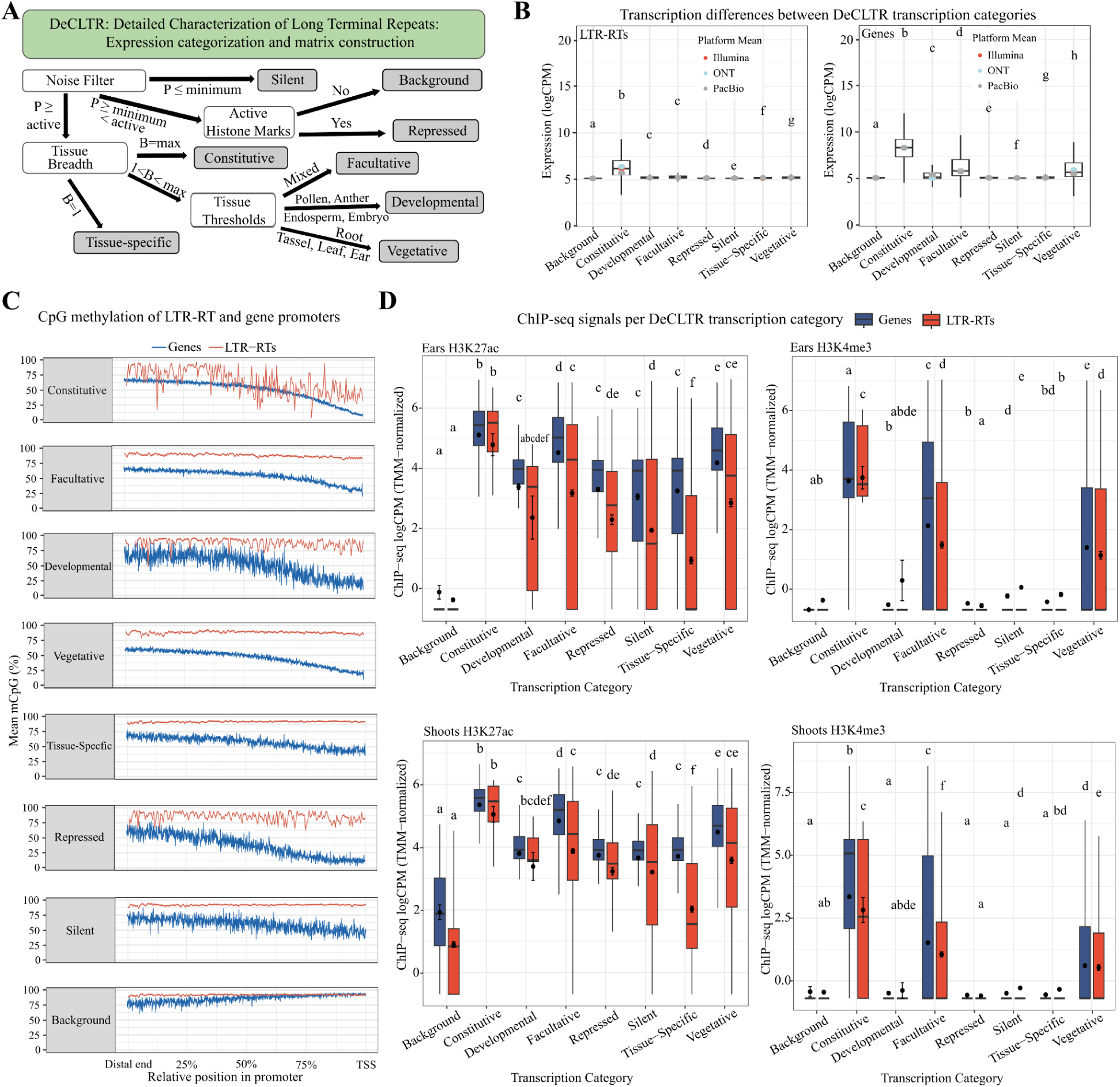
LTR-RTs and genes share similar expression and epigenetic profiles among transcriptional categories. **A)** Threshold criteria for transcriptional activity categorization using DeCLTR, which calculates transcription probability *P* in combination of DNA methylation and active histone marks across tissue breadth *B* for final categorization. **B)** Transcript abundances of LTR-RTs (left) and genes (right) between transcription categories defined by DeCLTR. Different letters between groups suggest statistical significance (Games-Howell test, *p* < 0.05). Colored points are the median transcript counts per sequencing platform. **C)** Mean per-base mCpG for DeCLTR categories that include 32,736 structurally intact LTR-RTs and 37,925 genes. The TSS of transcribed LTR-RTs were imputed to those non-transcribed family members for a total of 237 unique families. LTR-RT promoter is defined as the 5*′* LTR-RT start to the TSS, while the gene promoter is defined as 1kb upstream of the TSS. **D)** Distribution of normalized ChIP-seq read counts in the promoter regions of transcribed genes and LTR-RTs separated by DeCLTR categories. Different letters suggest statistical significance (Games-Howell test, *p* < 0.05). Error bars depict standard error.

Using DeCLTR, genomic loci with high transcript support in all tissue groups were sorted into the “Constitutive” category. Constitutive loci had the lowest levels of CpG methylation in their promoter regions and carried abundant active histone marks (**Fig. 3C,D**; **Fig. 4A,B; Fig. S2**). Similar to constitutive genes, Constitutive LTR-RTs had more abundant transcripts (**Fig. 3B**) and consistently more abundant active histone marks than other LTR-RT loci (**Fig. 3D**). Genes and LTR-RTs without sufficient transcript support were separated into “Silent” or “Repressed” categories, where the Repressed category was distinguished by CpG hypomethylation and the active histone marks H3K4me3 and H3K27ac (**Fig. 3A**). A Games-Howell test found “Developmental” and “Vegetative” LTR-RTs did not have statistically distinct H3K27ac profiles from Repressed elements (*p* > 0.05; **Fig. 3D**). When combined with CpG hypomethylation, this indicates Repressed LTR-RTs may be transcriptionally poised with RNAP-II, a noted effect of H3K27ac^50^.

**Figure 4.**
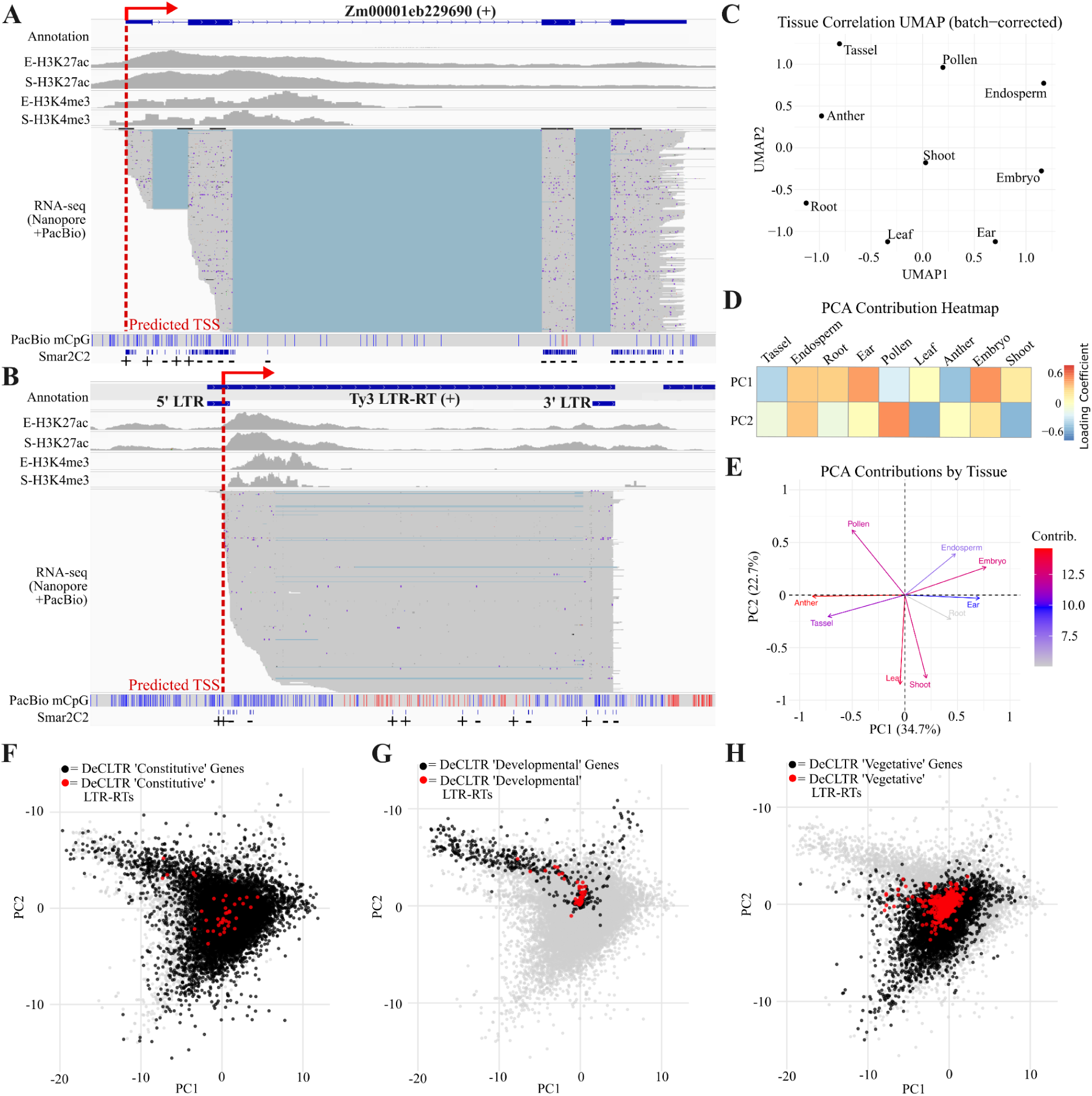
LTR-RTs have gene-like architecture and transcriptional clustering. Examples of **A)** a Constitutive gene and **B)** a Constitutive LTR-RT showed complementary and similar omics evidence. H3K27ac and H3K4me3 are shown for ears (E) and shoots (S). Blue bars and red bars on the PacBio mCpG track represent unmethylated and methylated cytosine, respectively. **C)** UMAP global transcription clustering between tissue groups used in DeCLTR. **D)** Stratification of tissue loading coefficients across PC1 and PC2 of the DeCLTR expression matrix. **E)** Tissue contributions to PC1 and PC2 shown by PCA biplot. Arrow direction and length indicate each tissue’s scaled loadings on the two components, while color reflects the overall contribution score to PC1 and PC2. **F, G, H)** PCA of all genes and LTR-RTs colored by DeCLTR activity groups of Constitutive **(F)**, Developmental **(G)**, and Vegetative **(H)**. Clusters correspond to tissue contributions in **E**.

Using transcriptomics data, Uniform Manifold Approximation and Projection (UMAP) separated tissues by regulatory state rather than organ identity (**Fig. 4C**). Principal Component Analysis (PCA) additionally showed differential contribution of each tissue to the principal components (**Fig. 4 D,E**). The PCA loading coefficients for PC1 and PC2 suggested that reproductive tissues of pollen and endosperm contributed most to PC2, while PC1 was driven by diverse tissues like ear, embryo, endosperm, and root (**Fig. 4D,E**). Similar clustering of genes and LTR-RTs was found in matching DeCLTR activity groups (**Fig. 4F-H**), and overlaying with the PCA biplot (**Fig. 4E**) showed Developmental-classified genes and LTR-RTs were appropriately concentrated with pollen, endosperm, and embryo (**Fig. 4D, E, G**). The Vegetative-classified elements clustered mostly away from the Developmental group, aligning with the arrows for tissues like leaf, shoot, root, and ear (**Fig. 4D, E, H**). Elements in the Constitutive group showed dispersion across all tissues (**Fig. 4E,F**). LTR-RTs in different DeCLTR categories appeared distinctly positioned throughout the tissue clusters, indicating their transcription is tissue-dependent (**Fig. 4F-H**). Importantly, LTR-RTs also showed varying levels of transcription, differential active histone marks, and DNA methylation in promoter regions among activity groups of Facultative, Tissue-Specific, Developmental, and Vegetative (**Fig. 3B-D**), suggesting context-dependent regulation similar to genes.

### Hub-like connectivity of co-expressed LTR-RTs and genes

Following analysis of broad transcription patterns, we were interested in the coexpression of LTR-RTs with genes. We performed a weighted-gene coexpression network analysis (WGCNA) of our multi-tissue transcriptomics dataset (**Table S1**) in R^52^. The analysis produced 76 coexpression modules. Each entry in the resulting heatmap (**Fig. 5A**) represents a module’s correlation with a tissue via its module eigengene (ME). There were 78 significantly correlated module-tissue pairs (Benjamini-Hochberg, FDR < 0.05), 60 of which contained LTR-RTs connected to genes (**Fig. 5A,B**). For this subset of LTR-RT-containing modules, we calculated module membership (kME) and gene significance (GS, the correlation of each element’s expression with a given tissue) to characterize the role of individual LTR-RTs within their respective coexpression networks. We then filtered networks by the most significant kME, GS, and topological connectivities (kME ≥ 0.75, kME *p* < 0.05, GS ≥ 0.5, GS *p* < 0.05) to more closely examine connections between LTR-RTs and genes (**Fig. 5C**).

**Figure 5.**
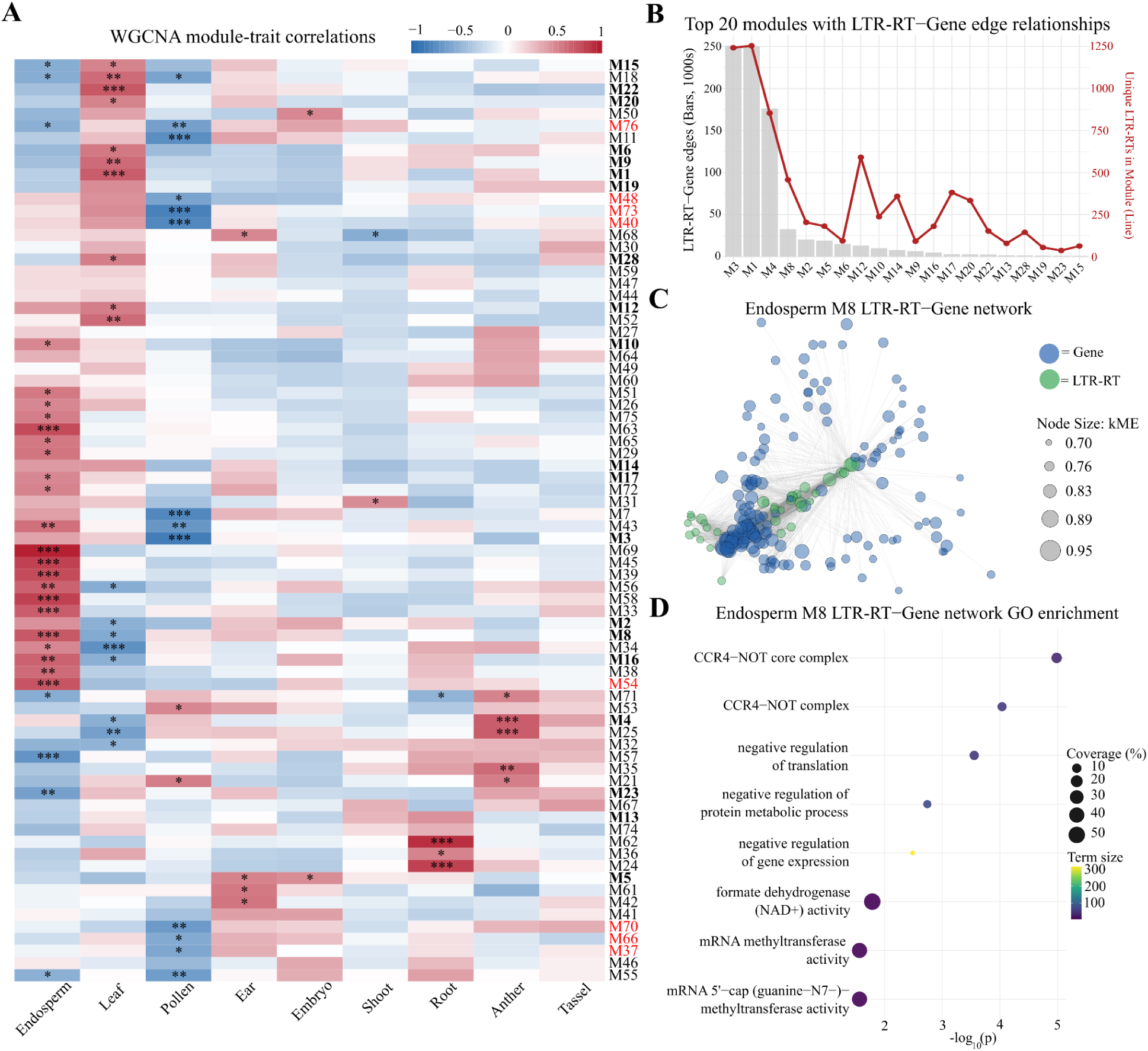
Weighted-gene coexpression analysis shows LTR-RTs with significant hub-like transcription. **A)** Module-trait correlation heatmap of coexpressed network modules in maize tissues. Modules that lack LTR-RT-gene connections are highlighted in red. The heatmap shows correlation -1 ≤ r ≤ 1 with significance levels: *, FDR ≤ 0.05; **, FDR ≤ 0.01; ***, FDR ≤ 0.001; **B)** Modules with top 20 high confidence LTR-RT-gene topological overlap measure (TOM) edges (bolded modules in **A**), and number of unique LTR-RTs in each network (red line). **C)** Visualization of the network in module M8 from endosperm with LTR-RTs (green) showing high edge weight connections (gray lines) to genes (blue). **D)** GO term enrichment of genes connected to LTR-RTs in the M8 endosperm module.

In LTR-RT-gene networks (LGNs), we were interested in functional enrichments that could imply LTR-RT activity in gene regulatory networks^53^. Of the 60 LGNs, 1,346 genes from 52 networks passed our filtering criteria. We conducted a gene ontology (GO) enrichment test using g:Profiler^54^ of those 1,346 genes to determine predominant cellular components, biological processes, and molecular functions in LGNs. Our analysis revealed 20/52 modules with enriched GO across leaf, anther, endosperm, pollen, ear, root, and shoot tissue. Endosperm contained the highest diversity of enriched networks: Module M8 was enriched for the CCR4-NOT core complex, the central component of cytoplasmic mRNA deadenylation and decay (**Fig. 5D**; **Table S4**); Module M29 was strongly enriched for rRNA processing, ribosome biogenesis, and nuclear exosome (RNase complex) activity, pointing to RNA surveillance machinery co-expressed with LTR-RTs; Module M33 was enriched for metabolic processes; and module M58 was enriched for the INO80 and SWI/SNF superfamily chromatin remodeling complexes (**Table S4**). The association of chromatin remodelers with LTR-RT–connected genes in endosperm is particularly relevant, as these complexes control nucleosome positioning and chromatin accessibility for genomic loci.

In the leaf, module M1 was enriched for photosynthesis and plastid localization, and module M28 showed enrichment for lysine N-methyltransferase activity, implicating protein methylation association with LTR-RT-coexpression. Modules M8, M16, and M34 shared enrichments for formate dehydrogenase (NAD+), glucose catabolism and aerobic respiration (processes anchored by phosphoglycerate mutase and phosphoglucomutase activity) across both endosperm and leaf but with reciprocal correlations (positive and negative, respectively; **Fig. 5A**; **Table S4**).

In the anther, module M4 was enriched for Rho protein signal transduction and enzyme inhibitor activity, while M35 showed enrichment for cell wall components. Root modules M24 and M36 were enriched for oxidoreductase and peroxidase activity including hydrogen peroxide metabolism and reactive oxygen species response, and plasma membrane localization, respectively. In pollen, module M21 was enriched for myo-inositol transport, and in ear, module M42 was enriched for RNA modification. A shoot-specific module, M31, was enriched for response to UV and blue light and plastid localization (**Table S4**). Collectively, these results revealed specialized and tissue-specific regulatory processes for module-trait associations involving LTR-RTs, with particular evidence for chromatin remodeling, RNA surveillance, metabolism, and post-transcriptional regulation.

Hub status in WGCNA is assigned to nodes/features most critical to their associated network based on kME, GS, topological overlap measure (TOM), and edge weight values. Interestingly, for some modules, we noted that LTR-RTs were assigned hub status over gene candidates, indicating their membership had a higher impact on network topology. We present two of these high-connectivity LTR-RT hubs as examples in **Fig. 6**. A hub-like *huck_AC216048_13250* LTR-RT element in M1 showed elevated transcript abundance in leaf and shoot tissues (**Fig. 6A**), and a single-copy LTR-RT insertion on chromosome 7 displayed elevated expression in endosperm and suppressed levels in leaf in the M8 module (**Fig. 6B**).

**Figure 6.**
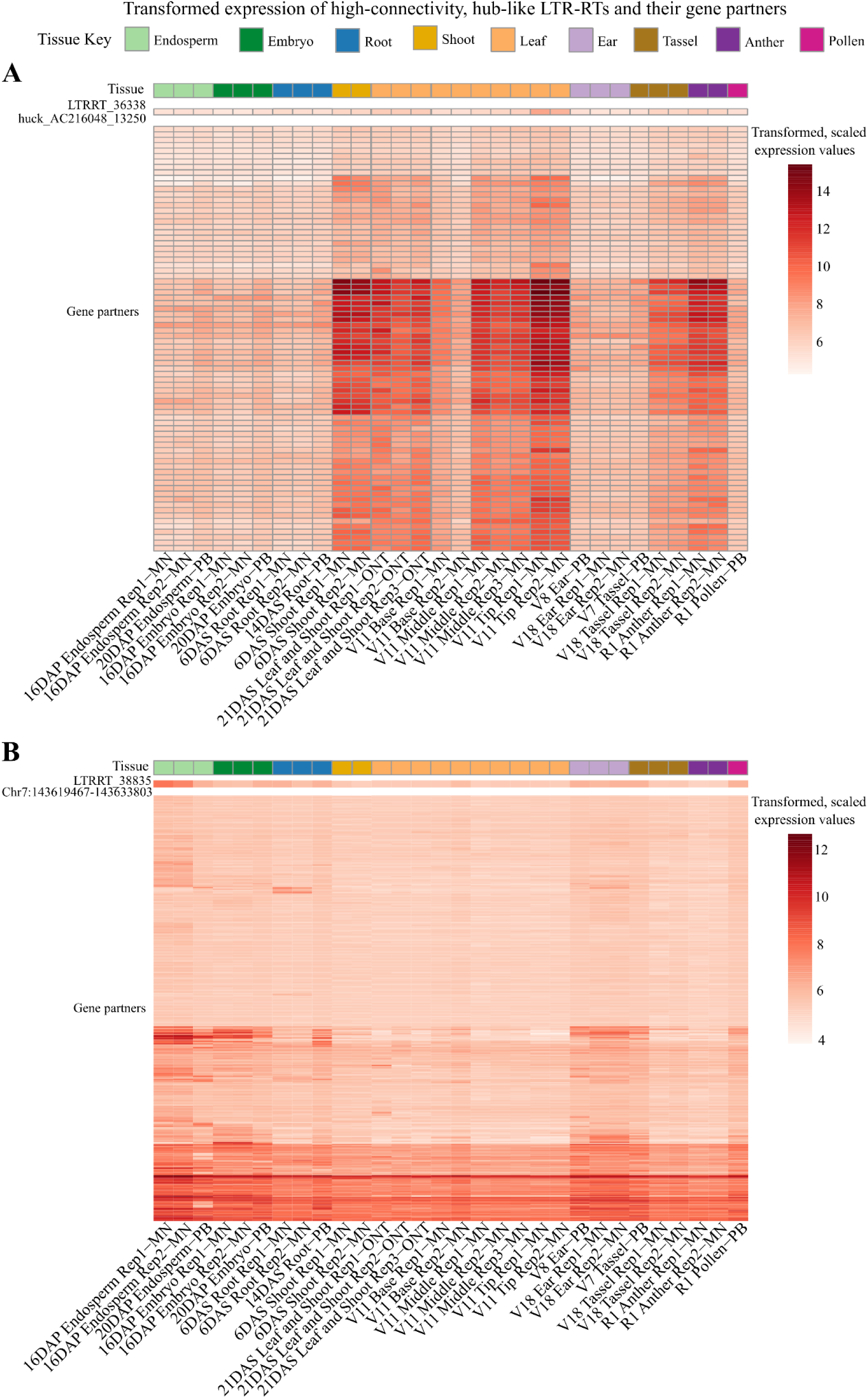
Examples of LTR-RTs displaying tissue specific expression patterns correlated with genes. Normalized expression of **A)** LTR-RT 36338 from the *huck_AC216048_13250* family in WGCNA module M1 and a single-copy LTR-RT 38835 on chromosome 7 in module M8 with gene partners from their respective modules. Gene partners: each row corresponds to the normalized expression levels of M1 genes (n = 78) and M8 genes (n = 474) connected to respective LTR-RT. DAP: day-after-pollination; DAS: day-after-sowing; MN: Illumina sequenced; ONT: Nanopore sequenced; PB: PacBio sequenced.

We examined family richness amongst transcribed LTR-RTs. First, we tested globally whether any LTR-RT families had high levels of intact elements transcribed and found 67 significantly enriched amongst all LTR-RT families (BH-adjusted, FDR ≤ 0.05; **Table S5**). When testing over-representation of LTR-RT families amongst each WGCNA module, we found 345 enriched families across 49 modules (BH-adjusted, FDR ≤ 0.05; **Table S6**). In addition, we observed that over 44% of single-copy LTR-RTs were transcribed and ∼29% appeared connected to genes in our WGCNA (**Table S7**), particularly in reproductive tissues like pollen, anther, and endosperm. Divergent expression signatures, association with enriched GO, and hub-like status in significant WGCNA modules suggests that LTR-RTs are deeply integrated into specialized, tissue-specific regulatory networks critical for biological procedures and functions.

## Discussion

Advances in long-read sequencing have improved TE studies by enabling full resolution of repetitive regions and precise transcript alignments^20,55^. LTR retrotransposons are a subclass of retrotransposons abundant in maize and other plants that have roles in regulating gene expression^28,56–59^, including as producers of lncRNA^19,31,60,61^ and other RNA species. Despite a conserved structural evolutionary history^13^, the ambiguous sequence composition of LTR U3 promoters obscures the mechanisms used to initiate transcription in the midst of sequence diversity, mutation, and host degradation^1,11^. This work takes advantage of the improved precision of long-read RNA sequencing to clearly identify the transcription start sites, and therefore delineating the U3 promoters of LTR-RTs displaying transcript abundance. Characterization of LTR-RT promoters and transcription is a critical first step to studying their discrete molecular roles in host genomes.

We found that long RNA-seq reads are reliable for determining TSS, a consensus primary site supported by ∼17% of reads aligned to genes and over 26% of those aligned to LTR-RTs. Our development and use of IsoClassifier to identify TSS also suggested that LTR-RTs predominantly produced transcripts from only the LTR region in our samples, and that the majority of these transcripts were spliced. The addition of PacBio Iso-seq reads supported this conclusion. Combined with transcript abundance from the non-coding LTRs, read support for canonical splice junctions and poly(A) tail presence supports post-transcriptional processing of LTR-RT transcripts to lncRNAs. It is well-documented that LTR-RTs and other TEs have insertional preference for gene intronic regions that can alter gene structure and function^1,62–65^, but to the authors’ knowledge, this is the first evidence suggesting that the LTRs of retrotransposons themselves contain introns relevant to their biology. Our results support LTR-RT’s major contribution of lncRNAs^66,67^ and shed light on their transcriptional processing for potential biological functions.

The variable length and composition of LTR regions previously made motif analysis difficult beyond broad generalizations. Defining the bounds of the U3 region made it possible for us to comprehensively explore the motif composition of 594 LTR-RT promoters from our combined ONT and PacBio RNA-seq analysis. We found similar core promoter motif composition in LTR-RTs compared to genes despite LTR-RTs’ higher mutation rates, more frequent epigenetic silencing, and a likely shorter average promoter lengths (i.e., LTR-RT 5’ start to TSS). The promoter similarities between LTR-RTs and genes are additionally surprising given the need for LTR-RTs to carry RNAP-II recruitment machinery with them during their transposition to avoid reliance on surrounding genomic contexts for transcription. Our long-read RNA-seq data revealed canonical core promoter motif profile in addition to the abundance of *de novo* motifs enriched in transcribed LTR-RTs, which provides resources for future LTR-RT sequencing experiments and annotation refinements.

The similarities in LTR and gene promoters were also reflected in the epigenetics data in **Fig. 3**. Silenced LTR-RTs displayed significantly high levels of DNA methylation as reported previously^22,68^. Repressed LTR-RTs showed distinct histone modification footprints and activation potentials compared with facultative LTR-RTs and other transcribed categories, likely due to similarities in silencing machinery but unique transcription priming (**Fig. 3C**).

The histone mark H3K4me3 at the 5′-end of genes is linked with active transcription, open chromatin, and regulatory hubs for essential gene functions^48,49^. Elevated levels of H3K4me3 overlap in some LTR-RTs imply they are not only affecting gene expression through random insertion but may be selected as regulators in a tissue-specific manner. The results from WGCNA also support this hypothesis. In leaf module M1 and endosperm module M8, for example, highly significant hub-like membership of LTR-RTs in these networks imply their participation in the tissue-specific transcriptional programs consistent with the enriched GO terms. Interestingly, though both the example LTR-RTs in **Fig. 6** had histone modifications and a hypomethylated CpG island, the *huck_AC216048_13250* element only connected to 78 genes while the single-copy LTR-RT in endosperm had over 400 edge partners. This could be a result of lessened genomic silencing in the germline, allowing increased LTR-RT transcription and potential partners. When combined with our data showing predominant transcription of spliced non-coding RNA from LTR regions, this supports a model wherein some LTR-RTs may have been domesticated for *trans*-regulation of gene expression in addition to known effects by *cis*-regulatory activity or as a result of insertion. The high WGCNA significance of single-copy LTR-RT insertions in reproductive tissues is unsurprising given newer insertions are generally more active in the germline. What is surprising, however, is that most of the transcription aligned to these copies appears non-coding and so could not be used for transposition. The behavior, function, and evolutionary impact of these single-copy LTR-RTs requires further study.

Since the 5′ LTR is identical to the 3′ LTR, both sequences contain a termination signal. It is hypothesized that 5′ LTR sequence composition suppresses termination through structural inhibition or protein binding, allowing RNAP-II to bypass the 5′ LTR poly(A) signal (PAS)^69,70^. We suspected that recent mutations in the 5′ LTR during transposition removed these capabilities and reinstated the 5′ PAS function, and similar mutations could also have allowed transcription to initiate from the 3′ LTR. The insertion time of LTR-RT estimated using Kmer2LTR^32^ supported this hypothesis, showing that loci producing predominantly lncRNA-like transcripts have an age distribution similar to the coding transcript-dominant loci (likely single-copy, new insertions). We also found that LTR-RTs expressing spliced LTR-contained transcripts are slightly younger than the non-spliced-producing loci, where the latter have average LTR age closer to silent elements.

The tools presented in the first half of this paper provide a method that can be used to identify the TSSs and transcript isoforms of transcribed LTR-RTs, the sequencing information of which can then be used for whole-genome analyses. We also addressed outstanding questions in the field of TE biology concerning transcription initiation of mobile retroelements and found evidence for introns and intronic splicing within LTR regions. Future study may focus on investigating the abundance of this splicing behavior in other eukaryotes, supported by the generalization of our pipelines. In addition to advancing the current dogmas of LTR-RT biology, the delineation of U3 promoters provides a unique opportunity for bioengineering. IsoClassifier and DeCLTR found distinct tissue-specific and developmentally-involved LTR-RT isoforms in maize, and can accomplish the same task in other model systems. If tissue-specific LTR-RT isoforms are identified, it is plausible that the U3 promoter sequences from those LTR loci could be isolated and used for molecular cloning to target a payload in an organism using an endogenous promoter. This would have the benefit of specificity without the side effects of overexpression sometimes observed when using exogenous and/or constitutive promoters^71^. It should also be noted that transcript abundance does not equate to active transcription, and so our conclusions are based on associations and trends of transcript abundance. Further evidence is required to conclude active transcription and to assess functional usage of lncRNA transcript isoforms. Collectively, this study presents a novel tool suite for characterizing transcribed LTR-RTs and studying their potential regulatory roles to enhance the field of TE biology and agricultural biotechnology.

## Data Availability

Raw Nanopore sequencing data generated in this work are available through the European Nucleotide Archive (ENA) BioProject ID PRJEB111639.

## Code Availability

All source code for the tools discussed in this work are publicly available on Github at https://github.com/cgooden-TE/LTRRT-Promoter-Suite. The version used in this study was archived on Zenodo: https://doi.org/10.5281/zenodo.19697466.

## Author Contributions

S.O. conceived and supervised the study. X.L. planted maize seedlings, prepared libraries, performed Nanopore sequencing and basecalling. C.G. developed custom bioinformatic pipelines, performed data analyses, created figures, and wrote the majority of the manuscript. I.W. processed the ChIP-seq data and called peaks. X.L. and I.W. wrote relevant method sections. All authors revised and agreed on the manuscript.

## Acknowledgements

1. G. was supported by NIH T32 GM141955. S.O. and X.L. were supported in part by the OSU STEM Education and JobsOhio award.

## Competing Interests

The authors declare no competing interests.

## Methods

### Plant material and RNA sequencing

Maize B73 seeds were soaked in 3% hydrogen peroxide (Fisher Scientific) overnight before sowing in pots. Three biological replicates were planted. Each replicate (n=12 plants) was grown in a chamber with 25°C, 16h light and 18°C, 8h dark at 60% of humidity. Three weeks after sowing, leaves were harvested followed by snap freezing in liquid nitrogen.

Total RNAs of maize tissues were extracted following the TRIzol Reagent (Invitrogen) protocol. In brief, ∼0.5g leaves were ground into fine powder in liquid nitrogen. The powder was suspended in 5 mL of TRIzol Reagent in a 15 mL tube at room temperature. After incubation, 1 mL chloroform was added and mixed thoroughly. The lysate was centrifuged, and the upper aqueous phase was transferred to a new 15 mL tube. The RNA was precipitated by centrifugation after mixing with isopropanol. After washing with 75% RNase-free ethanol, the RNA pellet was resuspended in RNase-free water.

The RNA-seq library construction was completed according to the manual of the ONT cDNA-PCR Barcoding Kit (SQK-PCB114.24). Firstly, RNA samples underwent quality control testing via Tapestation analysis (Agilent). Only samples with a RIN (RNA integrity number) number above 7.5 were used for subsequent processes. For each qualified sample, 500ng total RNA was taken as input for library construction. Each sample was mixed with cDNA RT Adapter and Annealing Buffer and incubated at 60°C for 5 mins. The ligation mix (NEBNext® Quick Ligation Reaction Buffer, T4 DNA Ligase, RNaseOUT) was then added to each tube. After ligation for 10 minutes at room temperature, the enzyme mix (Lambda Exonuclease, Uracil-Specific Excision Reagent) was added following incubation for 5 minutes at 37°C. Each sample was extracted by RNase-free XP beads (Beckman Coulter) and eluted into a clean Eppendorf tube. The RT Primer and dNTPs (10 mM) were added into each sample and incubated for 5 minutes at room temperature. Then, the reverse transcription mix (Maxima H Minus 5x RT Buffer, RNaseOUT, and Strand Switching Primer II) was mixed with the previous sample solution and incubated at 42°C for 2 minutes. Afterwards, 1 µl of Maxima H Minus Reverse Transcriptase was added to each tube for reverse transcription reaction in a thermal cycler. The reverse-transcribed samples were barcoded respectively during 14 cycles. All the amplified cDNA barcoded samples were pooled and ligated with an adapter in one cDNA sequencing library. Finally, the cDNA library was loaded in a PromethION R10.4 flow cell and sequenced in a Nanopore P2-solo device (Oxford Nanopore Technologies).

### Nanopore sequence processing and alignment

Nanopore raw cDNA reads were basecalled and demultiplexed using Dorado (v0.5.2+7969fab) with model ‘dna_r10.4.1_e8.2_400bps_sup@v4.3.0’ and option ‘--no-trim’ to retain primer and adapter sequences. Nanopore’s Pychopper v2.7.10 (RRID:SCR_018966) was used to orient and trim full-length reads using the “-k PCB114” option to match the cDNA Barcoding Kit (SQK-PCB114.24) used in the library preparation step. We developed a custom pipeline called “AccuMap” on Python 3.10.14 to retain strand orientations and pass reads to Cutadapt^24^ (v4.2) for homopolymer removal using “-a A{15} -g T{15}” parameters to remove poly(A) and poly(T) tails from the reads. The script then aligned reads to the B73v5 maize genome using splice-aware options “-ax splice -uf -k14 -secondary=no -G 20000” in minimap2 (v2.28-r1209) and created a custom BAM file that included the strand orientations as a tag. Rarefaction evaluation was performed using AlignQC^27^ (v2.0.5) provided the B73v5 genome and Nanopore transcriptome, and the results were plotted in R. Alignment statistics for LTR-RTs and genes were determined using featureCounts^72^ (v2.1.1), and alignments were visualized using the Integrative Genomics Viewer (IGV)^73^.

### ChIPseq peak calling

ChIP-seq datasets for H3K27ac and H3K4me3 modifications in maize immature ear and shoot tissues were obtained from NCBI BioProject PRJNA487471 (ref ^47^). Raw sequencing data (SRA format) were downloaded using parallel-fastq-dump (RRID:SCR_024150; v0.6.7) with --split-files and --gzip options. The B73v5 reference genome (Zm-B73-REFERENCE-NAM-5)^26^ and corresponding gene and TE annotations were downloaded from MaizeGDB. Gene and TE annotation files were filtered to remove noninformative or redundant features (exons, mRNAs, long terminal repeats, target site duplications, and repeat region annotations) to generate simplified feature sets for downstream intersection analyses. Reads were aligned to the B73 reference genome using Bowtie2 (ref ^74^)(v2.4.1) with default parameters. The resulting SAM files were converted to BAM format, sorted, and indexed using samtools^75^ (v1.9). Peaks were called using MACS3 (ref ^76^; v3.0.0a6) with a *p*-value threshold of 10e-5.

### DNA methylation calling

Long-read genomic sequencing datasets of B73 young leaves were obtained from the NCBI BioProject PRJNA627939 (ref ^77^). Raw sequencing reads were aligned to the B73v5 reference genome using pbmm2^78^ (v1.9.0) with mapping quality of 60 to eliminate false alignments. The resulting BAM files were merged into a single BAM file using samtools^75^ merge (v1.15.1). Methylation probability of CpG loci was generated using the machine-learning model and the reference mode provided by the pb-CpG-tools^79^ (v3.0.0). CpG loci with read coverage lower than 4 or higher than 52 (the mean coverage plus five times of the standard deviation) were determined unreliable and filtered out. The BSgenome^80^ (v1.66.3) and MethylSeekR^81^ (v1.38.0) packages were used to identify unmethylated regions (UMRs) from CpG methylation results using minimal coverage of 5, low-methylation cutoff of 50%, minimal number of 6 CpG loci for candidate UMR assignment, and 3 CpG loci for smoothing. Finally, candidate UMRs with average CpG methylation levels lower than 20% were retained as UMRs and used in this study.

### Smar2C2 and CAGE reprocessing

The original Smar2C2 results were presented using gene annotations^33^. It was necessary to reproduce the results for LTR-RTs in this study. To do so, reads from B73 maize shoot Smar2C2 sequencing were obtained from NCBI project PRJNA809198. We replicated the processing steps detailed in the Smar2C2 methods^82^. In brief, the reads were processed with Cutadapt^24^ v4.7 to remove the Smar2C2 TSS sequencing adapter, with umitools^83^ v1.1.0 to remove UMIs. The remaining paired-end reads were aligned to the B73v5 genome with STAR^84^ (v2.7.11b) to produce a single alignment file in BAM format. This alignment was used as input to TSRchitect^85^ (v.1.17.3) with default settings as done previous^82^, and the custom scripts in the Smar2C2 GitHub repository (https://github.com/aem11309/smar2C2) were modified to process the TSS and TSR outputs until we produced a discrete TSS set specific to our reference annotations for LTR-RTs. This set of TSS, containing the chromosome and start coordinate, were used as orthogonal evidence in the DeCLTR matrix (see below) for omics overlap and enrichment with IsoClassifier TSS predictions.

The CAGE TSS data were originally published using the B73v4 genome^34^. We obtained the B73v5 version from the USDA public dropbox collection (https://ars-usda.app.box.com/v/maizegdb-public/folder/164819527502) deposited by the original authors. We intersected the root and shoot dTSS files with our LTR-RT and gene annotation using BEDTools intersect^86^ (v2.26.0), then combined these into the DeCLTR matrix (see below) similarly to the Smar2C2 TSS to find overlap with predictions and for enrichment testing. We retained the TSS shape (broad or sharp) determined in the original CAGE experiment for additional characterization of LTR-RT promoters.

### Smar2C2 enrichment/depletion test

We tested whether Smar2C2 sites are non-randomly distributed with respect to transcription start sites (TSSs) of genes and LTR-RTs using a strand-aware, chromosome-preserving permutation approach. Each Smar2C2 site was represented as a 201 bp window centered on its genomic position and classified by overlap with either gene TSSs, LTR TSSs, intergenic regions, or regions of annotated elements outside our restrictions. We compared the observed proportions of Smar2C2 sites in each category against null expectations generated from 999 random permutations where 201 bp windows were randomly selected along the same chromosome and strand. For each permutation, we recalculated the proportions of overlaps and constructed an empirical null distribution. Observed enrichments or depletions were quantified as the ratio of the observed proportion to the permutation mean (fold-change = Observed / Expected). Empirical p-values were determined by the frequency with which permuted proportions exceeded (for enrichment) or fell below (for depletion) the observed proportion of Smar2C2 overlaps. Using this approach, we saw significant enrichment for Smar2C2 overlaps with gene and LTR TSSs compared to background and a slight depletion of sites in intergenic regions.

### PacBio IsoSeq processing and alignment

PacBio IsoSeq data from B73 was obtained from the NCBI BioProject PRJNA10769 (ref ^21^). Data from different sequencing runs were combined based on tissue type: root, ear, tassel, pollen, embryo, and endosperm. Raw reads were aligned to the B73v5 genome using minimap2 with splice aware parameters (-ax splice:hq -uf).

### Illumina NextSeq550 processing and alignment

Sequencing data from 10 maize tissues (embryo, endosperm, root, shoot, anther, leaf base, leaf middle, leaf tip, ear, and tassel) for B73 maize was obtained from the nested association mapping (NAM) parental sequencing project repository under NCBI BioProject PRJEB35943 (ref ^26^). Raw reads were aligned to the B73v5 genome using minimap2 in short-read mode (-ax sr), discarding secondary alignments. Read alignments intersected with genes and LTR-RTs using BEDtools intersect -c were counted intersections at each locus. Counts per locus per sample were bound into a master file via feature ID (TEs or genes) and merged with the DeCLTR matrix.

### Identifying transcribed LTR-RTs, corresponding U3 regions, and classifying isoforms

The “IsoClassifier” pipeline was developed for 1) finding real transcription events initiated from LTR-RTs, 2) identifying TSS candidates for genes and TEs, 3) using the primary TSS to delineate the U3 region of LTR-RTs, and 4) using the TSS and stranded reads to classify transcriptional activity into different isoform categories (**Figure 1**). IsoClassifier preferentially takes the alignment outputs of AccuMap or other long read BAM alignments and genome annotations of genes and LTR-RTs in GFF format. LTR-RTs nested inside other LTR-RTs and genes were removed to prevent false classifications. We used genomic coordinates of alignments and CIGAR information to determine introns and authentic primary alignments. A minimum mapping quality of 30 permissively filtered lowest quality reads, then we used strand flags native to minimap2 and strand tags added by PyChopper to further filter alignments, keeping those transcripts transcribed in the same genomic direction as the annotation. Final outputs included separate files for LTR transcript isoforms, primary and secondary LTR TSS counts, primary and secondary gene TSS counts, and two custom files summarizing TSS density. The aligned PacBio and Nanopore transcripts were unified and then classified by IsoClassifier.

The LTR primary/secondary TSS ID file was used as input for “U3_Seq_Extractor,” a small script that uses the B73 reference genome to determine ranges and sequences of U3 regions and LTRs based on TSS, strand, and annotation coordinates. The gene primary/secondary TSS ID file is used in a similar script which extracts the sequences of 1000 bp-ranges upstream of the TSS as the promoter region. The sequence files produced here are used as input for WindowScrubber to perform motif analysis.

### Motif analysis

Motif comparison of gene and LTR-RT promoters were completed with STREME^40^, AME^87^, and FIMO^44^ with selected sequences. Novel motifs for LTR-RT promoters were identified using full LTR sequences input to STREME in the MEME Suite. We compared enrichment of motifs in transcribed and silent LTR-RTs utilizing the STREME differential mode combined with AME, and tested positional distribution using FIMO.

In addition to utilizing tools in the MEME Suite, we developed WindowScrubber to perform motif analysis on extracted promoter sequences of both genes and LTR-RTs. This allowed more unique search windows for motif identification anchored on our TSS predictions. WindowScrubber combined sliding window and position weight matrix (PWM) approaches to identify predefined motifs within known ranges relative to the TSS in monocots^35^. The default ranges are as follows: TATA, -100 bp∼TSS+8 bp; Secondary TATA, TSS∼+100 bp; Initiator, ±10 bp at TSS; CCAAT, -460 bp∼-140 bp; Y-patch, -100 bp∼TSS; Downstream Promoter Element (DPE), TSS∼+50 bp.

The PWM approach determined the probability of each base at a position in our defined window, assigning a score to matches based on position-specific weights compared to background probabilities as the window slid through the specified range. We modeled core promoter motifs as PWMs where each column contains base probabilities at each position for A, C, G, and T according to consensus sequences as follows: TATA box, [TATAWAWR]; Initiator, [YTCANYY]; CCAAT box, [CCAAT]; Y-patch, alternating Y*8; Downstream Promoter Element (DPE), [RGWCGTG], in which W, R, Y, and N represent A/T, A/G, C/T, and A/G/C/T, respectively. Default PWMs were defined from “consensus” columns (strongly favoring the consensus base at each position), dual-base columns (two equally favored bases, ex. ‘W’), optionally blended with a uniform background to weaken tail positions as used for the last four bases of the TATA box. Users can optionally supply additional or alternative PWMs via a JSON file, which are merged with the defaults.

For scoring, each PWM is converted to a log-likelihood ratio (LLR) matrix using background nucleotide distributions, 0.25 by default. The LLR of each basepair is computed by

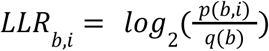

where *p(b,i)* is the PWM probability for base *b* at position *i* and *q(b)* is the background frequency. The sequences of each element were oriented according to strand and TSS, and scans were restricted to windows relative to the TSS based on previous reports in monocots^35^. For each motif, WindowScrubber applied a 1-bp–step sliding window of motif length across the specified range. For every window, motif score was computed as the sum of per-position LLRs. All windows with scores above the motif-specific scan threshold were recorded. For each motif PWM, a “perfect” score was computed as the sum of best-possible LLRs at each base position. Per-position score drops (difference between best and second-best base) were used to convert allowable mismatches *m* into a threshold. The “worst-case up to *m*-mismatches anywhere” cutoff was calculated by subtracting the largest drops from the maximum score. Mismatch allowances were set to zero by default so the strict mismatch-based thresholds equaled the perfect PWM score, but the scan and storage of motif hits used a lower, relative-score-based threshold, which is 85% a PWM’s dynamic range ± a 1.5-bit slack to maximize retention.

The threshold used for scanning motifs was permissive by default to allow maximum storage for later querying. The minimum possible LLR score of a motif is computed by summing the lowest LLR among A/C/G/T at each motif position, which is independent of mismatch settings and is computed to define the relative-score threshold. The relative score for each PWM is computed as

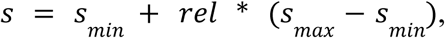

where *s* is the score and *rel* is a normalized value representing the possible score from 0.0-1.0, 0.85 by default. From this relative score, a slack of 1.5 LLR units was subtracted but floored at 0.0 to prevent negative motif matches (anti-motifs).

Finally, for each motif, an empirical null distribution of scores was generated using Monte Carlo sampling of random background k-mers and scoring them with the PWM. We used this score distribution to 1) convert each observed motif score for a match to a right-tailed *p*-value for probability under background, 2) to derive an empirical significance cutoff for p-values against a given Type-I error α (1 x 10^-4^ by default, following MEME Suite conventions^40,44^), and 3) to compute a percentile score (default = 95^th^ of observed hits) for each motif.

All motif hits above the threshold were written to a SQLite database with coordinates relative to the element and genome, distance to TSS, membership in TA-rich regions, motif sequence, LLR score, empirical *p*-value, and threshold values, enabling downstream selection of the “best” motif instances by score, distance, or significance. WindowScrubber is available in our GitHub repository (https://github.com/cgooden-TE/LTRRT-Promoter-Suite) and has an accompanying script “Query_WSDB.py” with some preset parameters used to summarize the resulting SQL database.

### Transcriptional activity categorization

Raw read count data from Nanopore (ONT), PacBio, and Illumina processing pipelines were incorporated with ChIPseq peaks and UMRs intersected with TE and gene annotations into a unified locus-by-sample matrix in R termed the **De**tailed **C**haracterization of **L**ong **T**erminal **R**epeats (DeCLTR) matrix. To reduce platform-specific background noise, we applied a two-stage segmented regression filtering approach. In the first stage, per-sample noise floors were estimated on raw (untransformed) count distributions. For each platform, we constructed a curve describing the number of loci with counts at or above an increasing integer threshold (1 to K, where K = 20 by default). A single-breakpoint segmented regression was fit to this curve using the R package *segmented*^88^. The estimated breakpoint transition from noise-dominated to signal-enriched was constrained to the interval [1, K] and rounded down to the nearest integer. For Illumina transcriptome data processing, thresholds were estimated independently for each sample. For PacBio data, thresholds were estimated per tissue by summing gene and LTR-RT counts per tissue. For ONT data, a single threshold was estimated from the sum of all merged control-tissue counts. These per-sample and per-tissue thresholds served as midpoints for the downstream sigmoid evidence function.

In the second stage, platform-level summary thresholds were estimated for filtering of transcribed loci. Per-locus read counts were computed by summing raw counts across all samples belonging to each platform, then transformed using log1p(*x*). The same segmented regression procedure was applied to these log1p-transformed totals to identify a platform-specific breakpoint in log-space. Loci with log1p total counts at or above the platform-specific breakpoint were retained as candidates for activity scoring. In addition, loci exhibiting chromatin evidence, either a ChIP-seq peak count exceeding its segmented-regression threshold or an unmethylated region (UMR) signal ≥ 0.1 (computed as 1 − mean methylation / 100), were retained regardless of expression level. Genes and LTR-RTs were not separated during thresholding; both feature types were pooled together at each stage so that noise floors reflect the combined expression landscape.

Next, samples were organized into nine biological tissue groups: leaf (including V11 Base, V11 Middle, and V11 Tip consolidated from B73 NAM), ear, shoot, root, anther, tassel, pollen, embryo, and endosperm. Within each tissue group and platform, locus-level expression was summarized as its median log1p(count) across replicate samples of that tissue–platform combination. For each locus, tissue group, and platform, activity evidence was calculated by comparing the median log1p expression to the corresponding noise-floor threshold. The difference (δ = median log1p expression − log1p threshold) was transformed into probabilistic evidence using a logistic sigmoid function:

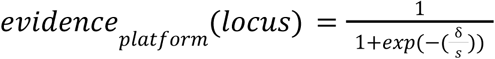

where s = 0.25 is a slope parameter controlling the sharpness of the transition. At the threshold δ = 0, evidence equals 0.5; expression well below the noise floor maps toward 0, and expression well above maps toward 1. For tissue groups sequenced by more than one platform, platform-specific evidence values were fused using a weighted average that ignores platforms lacking data for a given locus (NA-aware):

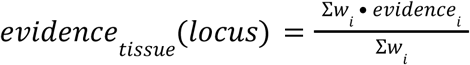

where the sum runs only over platforms *i* with finite evidence. Platform weights were set to *w*_Illumina_ = 1.0 and *w*_PacBio_ = 1.0, with *w*_ONT_ = 0.7 to reduce the influence of ONT on leaf-tissue activity calls where ONT was the only long-read platform available. These weights and the sigmoid slope were used to filter across all loci without per-feature-type tuning.

### Dimension reduction analyses

We transformed raw transcription counts into a standardized matrix to reduce platform-specific biases and facilitate comparative analysis across tissues. Raw count matrices from Illumina, PacBio, and ONT platforms were coerced into a starting matrix with padding for any missing features. We retained features with at least one count in at least one sample to ensure the inclusion of tissue-specific elements while removing completely uninformative features. Rather than a simple log-transformation, we employed a Variance Stabilizing Transformation (VST)^89^. We first calculated Trimmed Mean of M-values (TMM) as normalization factors using *edgeR*^90^ to determine effective library sizes. These were converted into size factors to parameterize a *DESeq2* model (design = ∼ platform + tissue). The resulting VST-transformed matrix stabilized the mean-variance relationship across the disparate sequencing depths of the three platforms. To integrate data across technologies, we used the *limma*^91^ package’s removeBatchEffect function, specifically protecting known tissue differences in the design matrix while removing the variance attributed to the sequencing platform. For tissue-level overview, we collapsed individual replicates into tissue groups using library-size weighted means based on TMM-adjusted effective library sizes. Subsequent Principal Component Analysis (PCA) and Uniform Manifold Approximation and Projection (UMAP) were completed using the adjusted expression unions.

To assess global relationship, each feature was normalized using standard deviations from the mean across tissue, measured by z-scores. PCA and UMAP (*n_neighbors* = 8, *min_dist* = 0.3) were performed on a transposed matrix using *base stats*^92^ (v4.3.2) and *uwot*^93^ (v0.2.3). We did not set a center or scale since our z-scored data was already corrected. UMAP analysis was done using a set.seed(1) for reproducibility. To examine the relationship between genes and LTR-RTs, instead of z-score normalization, we applied row-centering to the adjusted expression matrix to eliminate baseline expression differences while preserving the relative amplitude (magnitude of change) for each locus. We performed PCA using the prcomp() function. To ensure that tissues with higher global variance did not disproportionately dominate the principal components, we applied unit-variance scaling to the tissues (columns) within the function (scale = TRUE). Specific subsets were highlighted based on DeCLTR "Activity" classifications to observe the distribution of functional groups. Collectively, this allowed analysis focusing on the covariance of tissue-specific expression patterns. The influence of individual tissues on the resulting components was assessed using variable contribution plots (arrow plots) and scree plots using *factoextra*^94^ (v1.0.7).

### Weighted Gene Co-expression Network Analysis

The batch-corrected VST count matrices ordered by tissue and associated metadata were exported for WGCNA^52^ on our Linux server. Our input data included 21DAS leaf and shoot data sequenced by ONT; 20DAP endosperm and embryo, 14DAS root, V7 tassel, V8 ear, and R1 pollen sequenced by PacBio; and 16DAP embryo, 16DAP endosperm, 6DAP shoot, 6DAP root, V11 leaf tips, middles, and bases, V18 ears and tassel, and R1 anther sequenced by Illumina (**Table S1**). Samples were set as rows and genes/LTR-RTs as columns, then we removed low quality features with goodSamplesGenes() from WGCNA v1.73. The top 50% most varying features by median absolute deviation were retained to reduce noise. The tissue of origin served as the categorical trait. We constructed a signed network with biweight midcorrelation (bicor) using maxPOutliers = 0.1 to reduce weight from outliers, then used WGCNA’s pickSoftThreshold() to select a data-informed soft-thresholding power. We constructed networks with that soft power, corType = “bicor”, maxPOutliers, networkType & TOMType = “signed”, mergeCutHeight = 0.25 to merge modules with strongly correlated eigengenes, pamRespectsDendro = TRUE, and the standard color labels (numericLabels = FALSE). Module eigengenes (MEs) were ordered, correlated with traits, and assigned corresponding p-values computed using Student’s t approximation. After the analysis, modules were renamed using numbers instead of colors, and p-values were adjusted for multiple testing using the Benjamini-Hochberg procedure. In the custom processing, we also classified edge types (LTR-RT to LTR-RT, LTR-RT to gene, gene to gene) and recalculated the topological overlap matrix (TOM) per module as a quality check. For specific traits, gene significance (GS) was defined as the bicor between each feature’s expression and the trait, and module membership (kME) as the bicor between each feature and each ME. Module-trait relationships were visualized using a heatmap of correlation coefficients with annotated FDR-based significances. We used RCy3^95^ (v2.22.1) to pass our constructed networks directly to Cytoscape^96^ (v3.10.3) for visualization and used a custom script that queries the g:Profiler^54^ API (URL: https://biit.cs.ut.ee/gprofiler/api/gost/profile/) to perform GO enrichment on significant networks of LTR-RTs and genes.

### Artificial intelligence usage

Large language models were used to facilitate code development and computational accelerations.

## Notes

https://github.com/cgooden-TE/LTRRT-Promoter-Suite

### Competing Interest Statement

The authors have declared no competing interest.

## References

1. x Wells, J. N. & Feschotte, C. A Field Guide to Eukaryotic Transposable Elements. Annu. Rev. Genet. 54, 539–561 (2020).

2. Senft, A. D. & Macfarlan, T. S. Transposable elements shape the evolution of mammalian development. Nat. Rev. Genet. 22, 691–711 (2021).

3. Borredá, C., Leduque, B., Colot, V. & Quadrana, L. Transposable element products, functions, and regulatory networks in Arabidopsis. 2024.04.02.587720 Preprint at 10.1101/2024.04.02.587720 (2024).

4. Zhang, L. et al. A high-quality apple genome assembly reveals the association of a retrotransposon and red fruit colour. Nat. Commun. 10, 1494 (2019).

5. Tian, Y. et al. Transposon insertions regulate genome-wide allele-specific expression and underpin flower colour variations in apple (Malus spp.). Plant Biotechnol. J. 20, 1285–1297 (2022).

6. Wei, L. & Cao, X. The effect of transposable elements on phenotypic variation: insights from plants to humans. Sci. China Life Sci. 59, 24–37 (2016).

7. Studer, A., Zhao, Q., Ross-Ibarra, J. & Doebley, J. Identification of a functional transposon insertion in the maize domestication gene tb1. Nat. Genet. 43, 1160–1163 (2011).

8. Butelli, E. et al. Retrotransposons Control Fruit-Specific, Cold-Dependent Accumulation of Anthocyanins in Blood Oranges. Plant Cell 24, 1242–1255 (2012).

9. Lin, R. et al. Transposase-Derived Transcription Factors Regulate Light Signaling in Arabidopsis. Science 318, 1302–1305 (2007).

10. Bubb, K. L. et al. The regulatory potential of transposable elements in maize. Nat. Plants 11, 1181–1192 (2025).

11. Ou, S. et al. Differences in activity and stability drive transposable element variation in tropical and temperate maize. Genome Res. 34, 1140–1153 (2024).

12. Waters, A. J. et al. Natural variation for gene expression responses to abiotic stress in maize. Plant J. 89, 706–717 (2017).

13. Benachenhou, F. et al. Conserved structure and inferred evolutionary history of long terminal repeats (LTRs). Mob. DNA 4, 5 (2013).

14. Hirsch, C. D. & Springer, N. M. Transposable element influences on gene expression in plants. Biochim. Biophys. Acta BBA - Gene Regul. Mech. 1860, 157–165 (2017).

15. D’Orso, I. The HIV-1 Transcriptional Program: From Initiation to Elongation Control. J. Mol. Biol. 437, 168690 (2025).

16. Duverger, A. et al. An AP-1 Binding Site in the Enhancer/Core Element of the HIV-1 Promoter Controls the Ability of HIV-1 To Establish Latent Infection. J. Virol. 87, 2264–2277 (2013).

17. Suñé, C. & García-Blanco, M. A. Sp1 transcription factor is required for in vitro basal and Tat-activated transcription from the human immunodeficiency virus type 1 long terminal repeat. J. Virol. 69, 6572–6576 (1995).

18. Lanciano, S. & Cristofari, G. Measuring and interpreting transposable element expression. Nat. Rev. Genet. 21, 721–736 (2020).

19. Kirov, I. et al. Nanopore RNA Sequencing Revealed Long Non-Coding and LTR Retrotransposon-Related RNAs Expressed at Early Stages of Triticale SEED Development. Plants 9, 1794 (2020).

20. Wang, Y., Zhao, Y., Bollas, A., Wang, Y. & Au, K. F. Nanopore sequencing technology, bioinformatics and applications. Nat. Biotechnol. 39, 1348–1365 (2021).

21. Wang, B. et al. Unveiling the complexity of the maize transcriptome by single-molecule long-read sequencing. Nat. Commun. 7, 11708 (2016).

22. Panda, K. & Slotkin, R. K. Long-Read cDNA Sequencing Enables a “Gene-Like” Transcript Annotation of Transposable Elements. Plant Cell 32, 2687–2698 (2020).

23. Oxford Nanopore Technologies. Pychopper. Github [https://github.com/epi2me-labs/pychopper] (2024).

24. Martin, M. Cutadapt removes adapter sequences from high-throughput sequencing reads. EMBnet.journal 17, 10–12 (2011).

25. Li, H. Minimap2: pairwise alignment for nucleotide sequences. Bioinformatics 34, 3094–3100 (2018).

26. Hufford, M. B. et al. De novo assembly, annotation, and comparative analysis of 26 diverse maize genomes. Science 373, 655–662 (2021).

27. Weirather J.L., de Cesare M., Wang Y. et al. Comprehensive comparison of Pacific Biosciences and Oxford Nanopore Technologies and their applications to transcriptome analysis [version 2; peer review: 2 approved]. F1000Research, 6:100 (10.12688/f1000research.10571.2) (2017).

28. Xu, W., Thieme, M. & Roulin, A. C. Natural Diversity of Heat-Induced Transcription of Retrotransposons in Arabidopsis thaliana. Genome Biol. Evol. 16, evae242 (2024).

29. Yadav, V. K., Jalmi, S. K., Tiwari, S. & Kerkar, S. Deciphering shared attributes of plant long non-coding RNAs through a comparative computational approach. Sci. Rep. 13, 15101 (2023).

30. Mattick, J. S. et al. Long non-coding RNAs: definitions, functions, challenges and recommendations. Nat. Rev. Mol. Cell Biol. 24, 430–447 (2023).

31. Lv, Y., Hu, F., Zhou, Y., Wu, F. & Gaut, B. S. Maize transposable elements contribute to long non-coding RNAs that are regulatory hubs for abiotic stress response. BMC Genomics 20, 864 (2019).

32. Benson, C. W., Chen, T.-H., Thomson, S. J., Deng, C. H. & Ou, S. PrinTE: A Forward Simulation Framework for Studying the Role of Transposable Elements in Genome Expansion and Contraction. 2025.06.04.657780 Preprint at 10.1101/2025.06.04.657780 (2025).

33. Murray, A., Mendieta, J. P., Vollmers, C. & Schmitz, R. J. Simple and accurate transcriptional start site identification using Smar2C2 and examination of conserved promoter features. Plant J. 112, 583–596 (2022).

34. Mejía-Guerra, M. K. et al. Core Promoter Plasticity Between Maize Tissues and Genotypes Contrasts with Predominance of Sharp Transcription Initiation Sites. Plant Cell 27, 3309–3320 (2015).

35. Brooks, E. G. et al. Plant Promoters and Terminators for High-Precision Bioengineering. BioDesign Res. 5, 0013 (2023).

36. Yamamoto, Y. Y. et al. Differentiation of core promoter architecture between plants and mammals revealed by LDSS analysis. Nucleic Acids Res. 35, 6219–6226 (2007).

37. Kadonaga, J. T. The DPE, a core promoter element for transcription by RNA polymerase II. Exp. Mol. Med. 34, 259–264 (2002).

38. Kutach, A. K. & Kadonaga, J. T. The Downstream Promoter Element DPE Appears To Be as Widely Used as the TATA Box in Drosophila Core Promoters. Mol. Cell. Biol. 20, 4754–4764 (2000).

39. Jores, T. et al. Synthetic promoter designs enabled by a comprehensive analysis of plant core promoters. Nat. Plants 7, 842–855 (2021).

40. Bailey, T. L. STREME: accurate and versatile sequence motif discovery. Bioinformatics 37, 2834–2840 (2021).

41. Laloum, T., Mita, S. D., Gamas, P., Baudin, M. & Niebel, A. CCAAT-box binding transcription factors in plants: Y so many? Trends Plant Sci. 18, 157–166 (2013).

42. Gupta, S., Stamatoyannopoulos, J. A., Bailey, T. L. & Noble, W. S. Quantifying similarity between motifs. Genome Biol. 8, R24 (2007).

43. Ovek Baydar, D., et al. JASPAR 2026: expansion of transcription factor binding profiles and integration of deep learning models. 10.1093/nar/gkaf1209.

44. Grant, C. E., Bailey, T. L. & Noble, W. S. FIMO: scanning for occurrences of a given motif. Bioinformatics 27, 1017–1018 (2011).

45. Argelaguet, R. et al. Multi-Omics Factor Analysis—a framework for unsupervised integration of multi-omics data sets. Mol. Syst. Biol. 14, e8124 (2018).

46. Rappoport, N. & Shamir, R. NEMO: cancer subtyping by integration of partial multi-omic data. Bioinformatics 35, 3348–3356 (2019).

47. Li, E. et al. Long-range interactions between proximal and distal regulatory regions in maize. Nat. Commun. 10, 2633 (2019).

48. Foroozani, M., Vandal, M. P. & Smith, A. P. H3K4 trimethylation dynamics impact diverse developmental and environmental responses in plants. Planta 253, 4 (2021).

49. Beacon, T. H. et al. The dynamic broad epigenetic (H3K4me3, H3K27ac) domain as a mark of essential genes. Clin. Epigenetics 13, 138 (2021).

50. Creyghton, M. P. et al. Histone H3K27ac separates active from poised enhancers and predicts developmental state. Proc. Natl. Acad. Sci. 107, 21931–21936 (2010).

51. Stefansson, O. A. et al. The correlation between CpG methylation and gene expression is driven by sequence variants. Nat. Genet. 56, 1624–1631 (2024).

52. Langfelder, P. & Horvath, S. WGCNA: an R package for weighted correlation network analysis. BMC Bioinformatics 9, 559 (2008).

53. Feng, X. et al. Weighted Gene Co-Expression Network Analysis Reveals Hub Genes Contributing to Fuzz Development in *Gossypium arboreum*. Genes 12, (2021).

54. Kolberg, L. et al. g:Profiler—interoperable web service for functional enrichment analysis and gene identifier mapping (2023 update). Nucleic Acids Res. 51, W207–W212 (2023).

55. Ou, S. & Jiang, N. LTR_retriever: A Highly Accurate and Sensitive Program for Identification of Long Terminal Repeat Retrotransposons. Plant Physiol. 176, 1410–1422 (2018).

56. Bubb, K. L. et al. The regulatory potential of transposable elements in maize. Nat. Plants 11, 1181–1192 (2025).

57. Cahn, J. et al. MaizeCODE reveals bi-directionally expressed enhancers that harbor molecular signatures of maize domestication. Nat. Commun. 15, 10854 (2024).

58. Cavrak, V. V. et al. How a Retrotransposon Exploits the Plant’s Heat Stress Response for Its Activation. PLoS Genet. 10, e1004115 (2014).

59. Deneweth, J., Van de Peer, Y. & Vermeirssen, V. Nearby transposable elements impact plant stress gene regulatory networks: a meta-analysis in A. thaliana and S. lycopersicum. BMC Genomics 23, 18 (2022).

60. Cho, J. Transposon-Derived Non-coding RNAs and Their Function in Plants. Front. Plant Sci. 9, (2018).

61. Ariel, F. D. & Manavella, P. A. When junk DNA turns functional: transposon-derived non-coding RNAs in plants. J. Exp. Bot. 72, 4132–4143 (2021).

62. Domínguez, M. et al. The impact of transposable elements on tomato diversity. Nat. Commun. 11, 4058 (2020).

63. Wang, L. et al. Somatic variations led to the selection of acidic and acidless orange cultivars. Nat. Plants 7, 954–965 (2021).

64. Rohilla, M. et al. Genome-wide identification and development of miniature inverted-repeat transposable elements and intron length polymorphic markers in tea plant (*Camellia sinensis*). Sci. Rep. 12, 16233 (2022).

65. Wang, H. et al. The double flower variant of yellowhorn is due to a LINE1 transposon-mediated insertion. Plant Physiol. 191, 1122–1137 (2023).

66. Lv, Y., Hu, F., Zhou, Y., Wu, F. & Gaut, B. S. Maize transposable elements contribute to long non-coding RNAs that are regulatory hubs for abiotic stress response. BMC Genomics 20, 864 (2019).

67. Yadav, V. K., Jalmi, S. K., Tiwari, S. & Kerkar, S. Deciphering shared attributes of plant long non-coding RNAs through a comparative computational approach. Sci. Rep. 13, 15101 (2023).

68. Liu, B. & Zhao, M. How transposable elements are recognized and epigenetically silenced in plants? Curr. Opin. Plant Biol. 75, 102428 (2023).

69. Guntaka, R. V. Transcription termination and polyadenylation in retroviruses. Microbiol. Rev. 57, 511–521 (1993).

70. Fogel, B. L., McNally, L. M. & McNally, M. T. Efficient polyadenylation of Rous sarcoma virus RNA requires the negative regulator of splicing element. Nucleic Acids Res. 30, 810–817 (2002).

71. Villao-Uzho, L., Chávez-Navarrete, T., Pacheco-Coello, R., Sánchez-Timm, E. & Santos-Ordóñez, E. Plant Promoters: Their Identification, Characterization, and Role in Gene Regulation. Genes 14, 1226 (2023).

72. Liao, Y., Smyth, G. K. & Shi, W. featureCounts: an efficient general purpose program for assigning sequence reads to genomic features. Bioinformatics 30, 923–930 (2014).

73. Robinson, J. T. et al. Integrative Genomics Viewer. Nat. Biotechnol. 29, 24–26 (2011).

74. Langmead, B. & Salzberg, S. L. Fast gapped-read alignment with Bowtie 2. Nat. Methods 9, 357–359 (2012).

75. Danecek, P. et al. Twelve years of SAMtools and BCFtools. GigaScience 10, giab008 (2021).

76. Zhang, Y. et al. Model-based Analysis of ChIP-Seq (MACS). Genome Biol. 9, R137 (2008).

77. Hon, T. et al. Highly accurate long-read HiFi sequencing data for five complex genomes. Sci. Data 7, 399 (2020).

78. Pacific Biosciences. pbmm2. Github [https://github.com/PacificBiosciences/pbmm2] (2026).

79. Pacific Biosciences. pb-CpG-tools. Github [https://github.com/PacificBiosciences/pb-CpG-tools] (2026).

80. Pagès, H. BSgenome: Software infrastructure for efficient representation of full genomes and their SNPs. DOI [doi:10.18129/B9.bioc.BSgenome]. R package version 1.80.0. (2025).

81. Burger, L., Gaidatzis, D., Schübeler, D. & Stadler, M. B. Identification of active regulatory regions from DNA methylation data. Nucleic Acids Res. 41, e155 (2013).

82. Murray, A., Vollmers, C. & Schmitz, R. J. Smar2C2: A Simple and Efficient Protocol for the Identification of Transcription Start Sites. Curr. Protoc. 3, e705 (2023).

83. Smith, T., Heger, A. & Sudbery, I. UMI-tools: modeling sequencing errors in Unique Molecular Identifiers to improve quantification accuracy. Genome Res. 27, 491–499 (2017).

84. Dobin, A. et al. STAR: ultrafast universal RNA-seq aligner. Bioinformatics 29, 15–21 (2013).

85. Raborn, R., Sridharan, K. & Brendel, V. TSRchitect: Promoter identification from large-scale TSS profiling data. DOI [doi:10.18129/B9.bioc.TSRchitect] (2017).

86. Quinlan, A. R. & Hall, I. M. BEDTools: a flexible suite of utilities for comparing genomic features. Bioinformatics 26, 841–842 (2010).

87. McLeay, R. C. & Bailey, T. L. Motif Enrichment Analysis: a unified framework and an evaluation on ChIP data. BMC Bioinformatics 11, 165 (2010).

88. Muggeo, V. M. R. Estimating regression models with unknown break-points. Stat. Med. 22, 3055–3071 (2003).

89. Anders, S. & Huber, W. Differential expression analysis for sequence count data. Genome Biol. 11, R106 (2010).

90. Robinson, M. D., McCarthy, D. J. & Smyth, G. K. edgeR: a Bioconductor package for differential expression analysis of digital gene expression data. Bioinformatics 26, 139–140 (2010).

91. Ritchie, M. E. et al. limma powers differential expression analyses for RNA-sequencing and microarray studies. Nucleic Acids Res. 43, e47 (2015).

92. R Core Team. R: A Language and Environment for Statistical Computing. R Foundation for Statistical Computing. (2023).

93. Melville, J. uwot: The Uniform Manifold Approximation and Projection (UMAP) Method for Dimensionality Reduction. (2025).

94. Kassambra, A. & Mundt, F. factoextra: Extract and Visualize the Results of Multivariate Data Analyses. (2020).

95. Gustavsen JA, Pai S, Isserlin R et al. RCy3: Network biology using Cytoscape from within R [version 3; peer review: 3 approved]. F1000Research, 8:1774. DOI [10.12688/f1000research.20887.3] (2019).

96. Shannon, P. et al. Cytoscape: A Software Environment for Integrated Models of Biomolecular Interaction Networks. Genome Res. 13, 2498–2504 (2003).

