## Supplemental Figures for "The gene-like promoter and transcription of LTR retrotransposons"

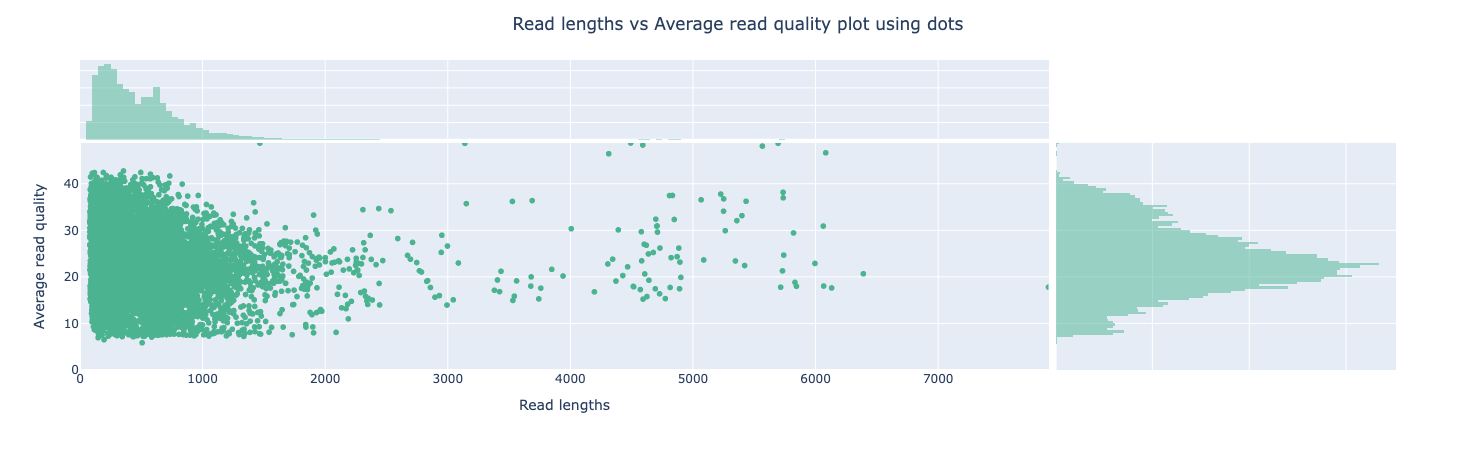

**Figure S1. Nanoplot distribution of read lengths versus read quality.** Average read quality plotted by read length after applying the initial filtering threshold in IsoClassifier (MapQ ≥ 30).

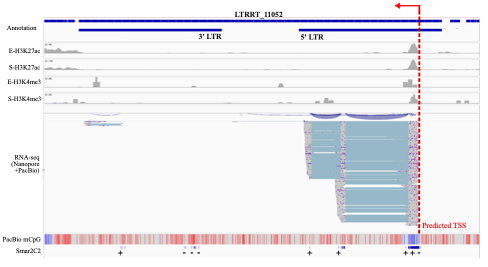

**Figure S2. Integrative Genomics Viewer (IGV) example of a spliced LTR-RT.** Histone modifications H3K27ac and H3K4me3 and CpG methylation support transcription. IsoClassifier TSS prediction exactly matches the Smar2C2-seq TSS evidence. Grey lines and blue lines on the RNA-seq track represent exons and introns, respectively. Blue bars and red bars on the PacBio mCpG track represent unmethylated and methylated cytosine, respectively.

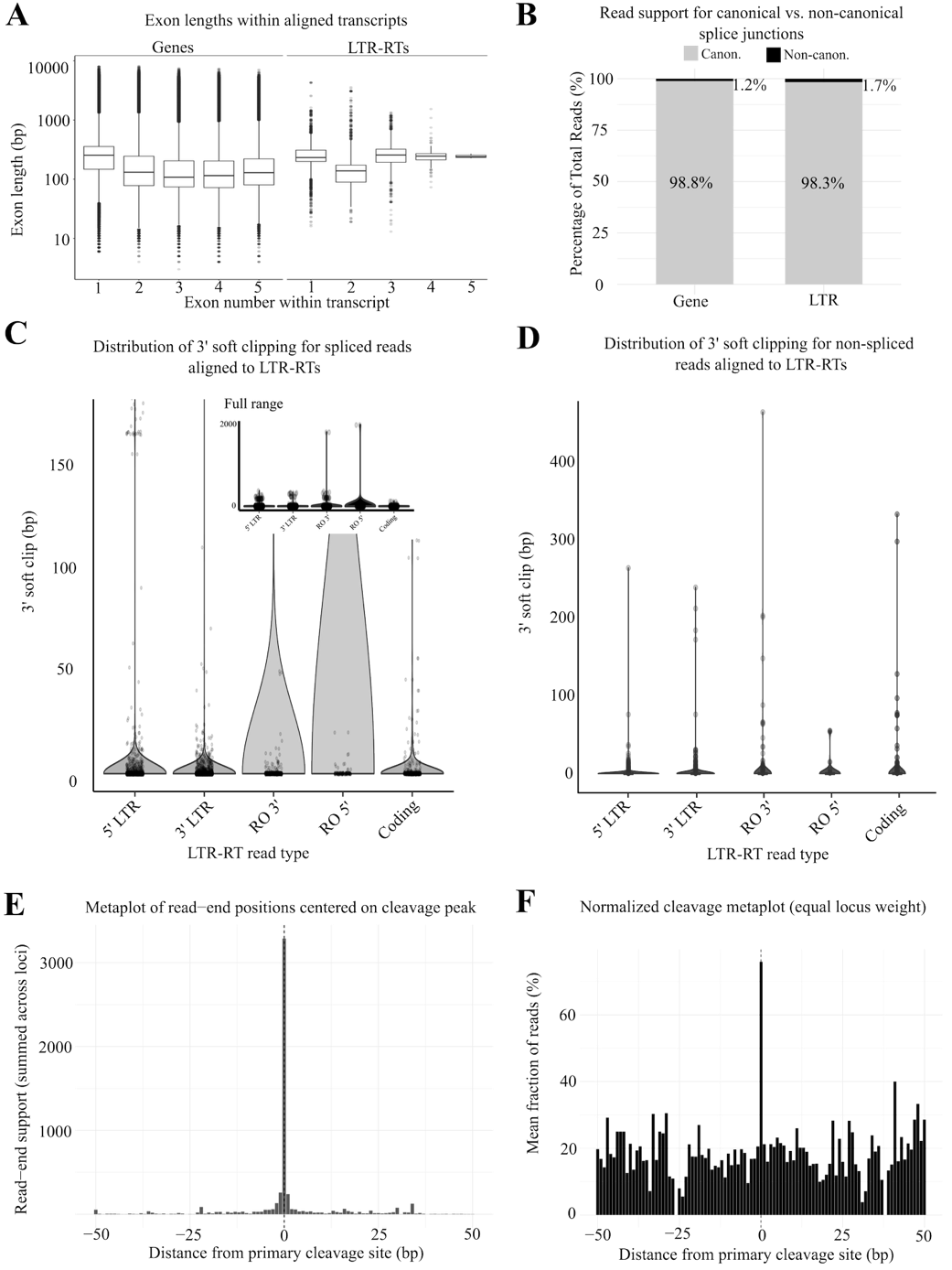

**Figure S3. Noncoding LTR-RT transcripts are spliced similarly to genes. A)** Length (bp, y-axis) distribution of the first five exons found in genes (left) and LTR-RTs (right). **B)** Percentage of total spliced reads that aligned to canonical splice junctions (gray) and non-canonical junctions (dark gray) showed near complete support for canonical junctions in both genes and LTR-RTs. **C)** Violin plot distributions of the length of sequences clipped from 3' read ends of spliced LTR-RT transcripts. The inset is the full data range, while the enlarged plot accounts for 99% of the data. **D)** Violin plot distributions of length of sequences clipped from 3' read ends of non-spliced LTR-RT transcripts. **E)** Distribution of read 3' end alignments across all transcribed LTR-RTs, with the x-axis representing the offset of each read’s 3’ end from the dominant alignment end (0; primary cleavage site). The y-axis is the read count summed across transcribed LTR-RTs. **F)** Distribution of normalized read 3' end alignments, where the y-axis is the mean of LTR-RT read ends *x* distance from the dominant alignment end. The sharp peaks at 0 indicate that read 3′ end aligned tightly on a single position per LTR-RT, with minimal dispersion into flanking bases.

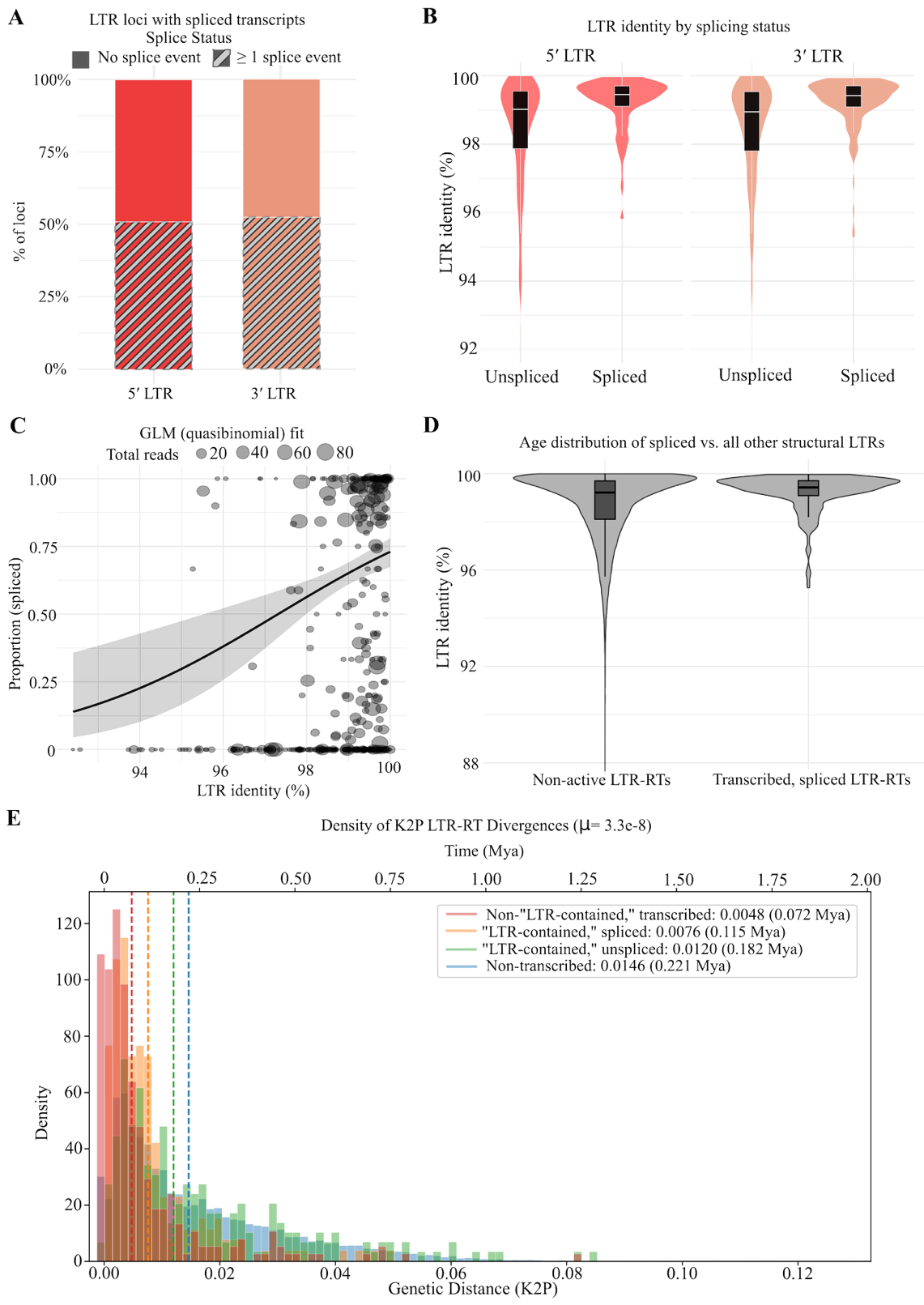

**Figure S4. Younger LTR-RTs produce predominantly spliced, non-coding transcripts. A)** Percentage of spliced transcripts from LTR-RTs that produced 5' or 3'-LTR-contained transcripts. **B)** LTR identity of LTR-RTs yielding spliced and unspliced 5' or 3'-LTR-contained transcripts. **C)** Quasibinomial generalized linear model (GLM) for LTR identity versus proportion of spliced transcripts. **D)** LTR identity of LTR-RTs yielding spliced transcripts versus all silent LTR-RTs in the genome showed that spliced LTR-RTs are younger. **E)** Estimation of LTR-RT locus age using Kmer2LTR. The x-axis shows the K2P genetic distance (bottom track) and the corresponding age in Mya (top track). Numbers in the legend are the K2P distances and age at density peaks. Kmer2LTR showed spliced LTR-RTs are younger than unspliced loci when using a mutation rate of µ = 3.3e-8 per bp per year for age estimations.

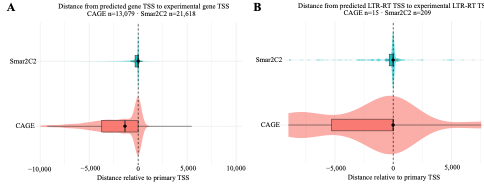

**Figure S5 Distribution of relative distance (bp) between IsoClassifier TSS prediction and closest TSS reported by Smar2C-seq2 and CAGE-seq in A) genes and B) LTR-RTs.**

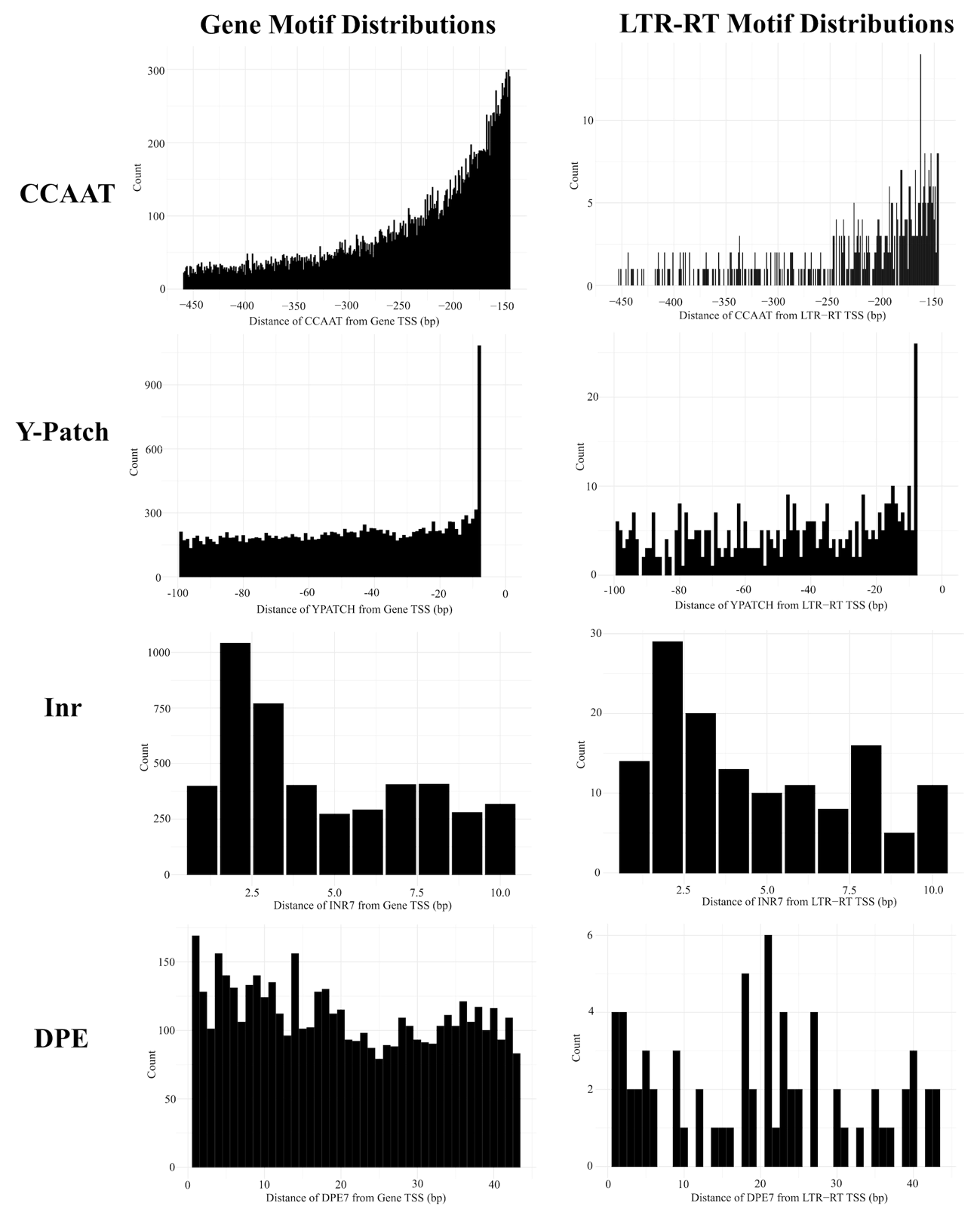

**Figure S6. Per-bp distribution of the highest-scoring core promoter motifs in LTR-RT and gene promoters.** Distributions of core promoter motifs relative to primary TSS for (top to bottom) CCAAT, pyrimidine- (Y) patch, initiator (Inr) sequence, and downstream promoter element (DPE) for genes (left) and LTR-RTs (right). Negative and positive values on the x-axis represent each motif’s canonical range upstream and downstream of the TSS, respectively. The y-axis represents the number of motif 5' ends at the corresponding x-coordinate within the range.

**Table S1. Genomic features overlapping with uniquely mapped raw ONT and PacBio reads, filtered (MapQ ≥ 30) ONT and PacBio reads, and Illumina short reads.**

**Table S2. Segmented regression breakpoints per sample used in the sigmoid function to determine expression probabilities.**

**Table S3. LTR-RT promoter motif annotations by TOMTOM using the non-redundant JASPAR CORE plants motif database with enrichments determined by STREME.**

**Table S4. Gene ontology enrichments in WGCNA modules containing LTR-RT and gene networks.**

**Table S5. Whole-genome enrichment of transcribed LTR-RT families based on one-sided Fisher’s exact test with Benjamini-Hochberg correction.**

**Table S6. Module-level enrichment of transcribed LTR-RT families based on one-sided Fisher’s exact test with Benjamini-Hochberg correction.**

**Table S7. 656 single-copy LTR-RTs found in 36WGCNA modules out of 1,590 transcribed and 3,599 total intact single-copy LTR-RTs genome-wide.**
